# Mosquito Ferritin: A Haem-Binding Iron Store Required for Egg Development and Targetable for Malaria Control

**DOI:** 10.64898/2025.12.01.691523

**Authors:** Chiging Tupe, Rahul Pasupareddy, Shruti Bhatt, Bharti Goyal, Ankit Kumar, Aashima Gupta, Amit lalwani, Rudra Nayan Das, Sujatha Sunil, Shailja Singh, Kailash C Pandey, Soumyananda Chakraborti

## Abstract

Iron metabolism is essential for both mosquito reproductive fitness and Plasmodium development within the vector, yet this axis remains insufficiently explored as a potential target for malaria control. Ferritin, the major iron-storage protein and a central regulator of iron homeostasis, has not previously been biochemically or functionally characterized in any hematophagous insect. Our computational and phylogenetic analyses reveal that mosquito ferritin represents a unique case, combining conserved iron-binding motifs with distinctive haem-coordinating residues that confer an exceptionally high haem-binding affinity exceeding that of bacterioferritin. In addition, through a screen of multiple iron chelators in Anopheles larvae, we identified potent candidates with potential utility for vector control. Together, these findings establish mosquito ferritin as an evolutionarily distinct and physiologically essential protein, and they highlight iron homeostasis as a promising dual-target axis for next-generation vector control strategies.

## Introduction

Malaria still remains one of the deadliest vectors borne disease globally. According to the global malaria report in WHO 2024 report ^1^, there have been increase in the malaria cases post pandemic. To exacerbate the situation, there have been recent reports of zoonotic malaria transmission from non-human primates (NHP) in South-East Asian countries ^2–4^. Eliminating NHP (non-human primate) malaria transmission has been particularly challenging due to the difficulty in reducing the parasite reservoir, *i.e.*, the NHPs themselves. Therefore, employing new malaria control strategies that focus either more on the vector (mosquitoes), or transmission stage of parasite in vector could be a more viable option. To achieve this, we need to understand mosquito biology in more details.

Conventional mosquito control strategies primarily rely on chemical insecticides such as pyrethroids, organochlorines, organophosphates, and carbamates, which target key metabolic pathways including voltage-gated sodium channels, GABA receptors, and acetylcholinesterase ^5–7^. However, no new classes of insecticides have been developed in the past three decades, leading to widespread resistance across mosquito populations. These challenges underscore the urgent need to find novel molecular targets for mosquito control. Emerging insights in genomics and proteomics are now revealing previously unrecognized, essential pathways that could serve as promising intervention points. Interestingly, despite its central role in the physiology of hematophagous insects, the iron metabolism pathway in mosquitoes remains largely unexplored. Notably, recent studies in bacteria have identified components of the iron homeostasis pathway as potential drug targets, suggesting a similar untapped opportunity in mosquitoes. Iron also acts as a cofactor for key stress-response genes such as Cytochrome P450, which has been identified as a major contributor to insecticide resistance ^8^. Beyond mosquitoes, other hematophagous vectors also heavily rely on iron metabolism for blood digestion, growth and development, as well as reproductive tissue development ^9^. However, unlike obligate blood feeders like tsetse flies and ticks, which require blood for survival and where both sexes feed on blood, mosquitoes are facultative blood feeders, with only females feeding on blood for egg development ^10,11^.

Within the intricate network of iron regulation and homeostasis, ferritin stands out as a central and evolutionarily conserved protein across nearly all life forms. Ferritin is a ubiquitous iron-storage molecule that assembles into a homo- or hetero-polymeric cage composed of 24 subunits. The heavy (H) chain possesses a ferroxidase center that catalyzes the oxidation of bioavailable Fe²□ into its stable Fe³□ form, while the light (L) chain facilitates iron nucleation and mineralization ^12^. Maintaining optimal Fe²□ levels in the cellular milieu is critical, excess Fe²□ can generate reactive oxygen species (ROS), leading to oxidative damage, whereas iron deficiency can impair Fe–S cluster biogenesis, thereby disrupting the function of numerous essential enzymes. To balance these opposing demands, nature has evolved ferritin as a dynamic iron reservoir, enabling safe storage and mobilization of iron during conditions of stress or metabolic demand. In mammals, ferritin typically forms a 24-mer composed of both H and L chains, whereas bacterial ferritins are usually homopolymeric, consisting of a single subunit type. Beyond its canonical role in iron storage, ferritin in insects also has been linked to contribute to several key physiological processes, including antioxidant defense, iron trafficking, immune regulation, and circadian rhythm maintenance ^13–17^.

Studies on insect ferritin and iron metabolism pathways remain limited. However, in a few individual studies on blood-feeding insects, ferritin has been shown to play essential roles beyond iron storage, particularly in supporting reproductive development and overall physiological fitness. RNA interference (RNAi)-mediated knockdown experiments in multiple insect species have demonstrated that reduced ferritin expression results in decreased fecundity, smaller egg size, and lower hatching success (**Table 1**). Structurally, insect ferritins are distinct, exhibiting tetrahedral symmetry rather than the conventional octahedral arrangement observed in vertebrates, with an H:L subunit ratio typically around 1:1 ^18^. Interestingly, both fully H-type and fully L-type 24-mer ferritin assemblies have been reported. Additionally, insect ferritins contain an N-terminal “leader peptide” that directs the nascent protein to the endoplasmic reticulum, where the signal sequence is cleaved prior to trafficking through the Golgi apparatus for secretion ^19^.

**Table 1.**
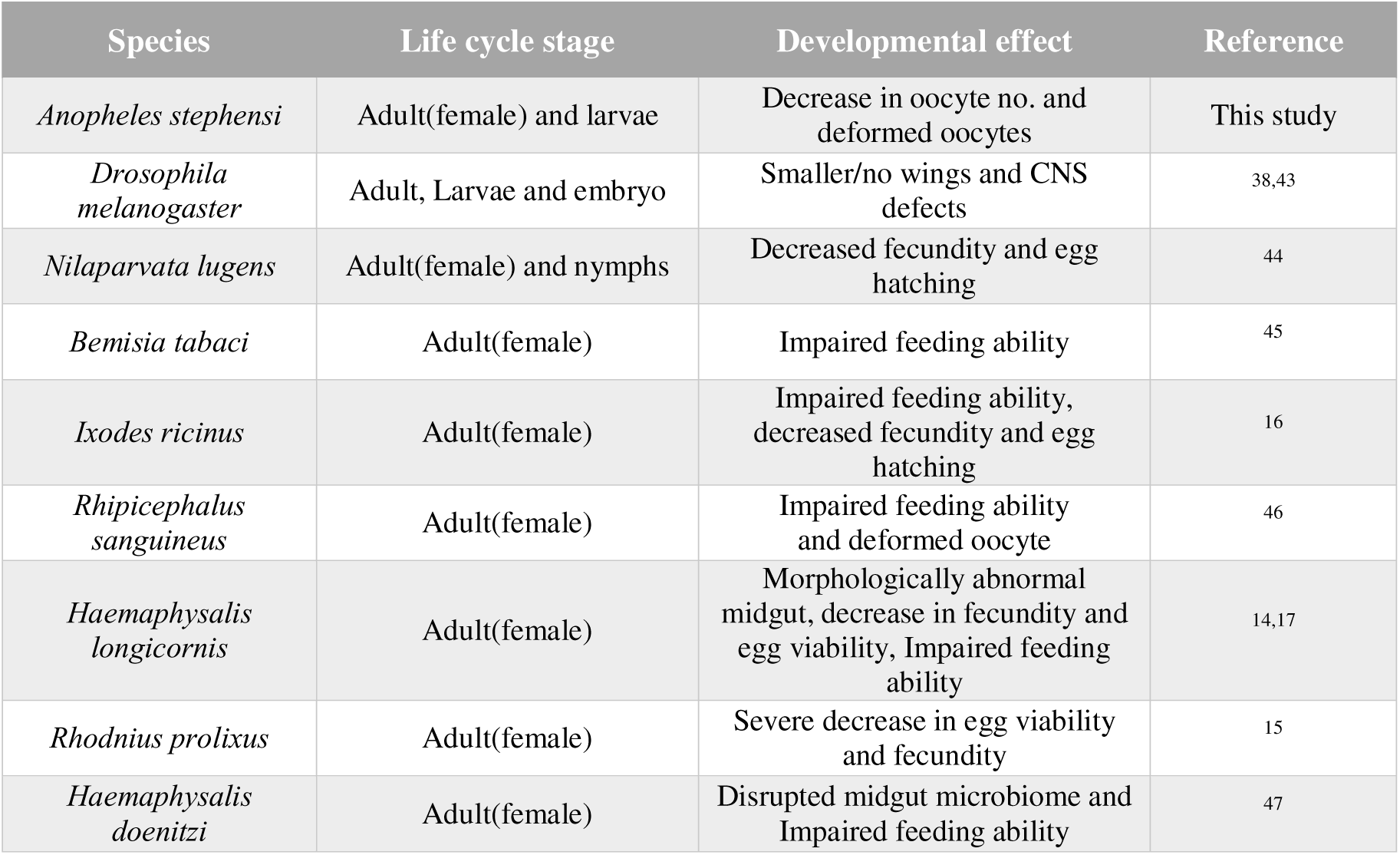
Comparison of RNAi-mediated knockdown effects of ferritin’s heavy chain homolog across different insect species.

**Table 2.** Comparative physicochemical properties of ferritin heavy chain homologs across different species. The data was generated using Protparam.

| Species | No.of amino acids | Sequence similarity (to *An.culicifacies*) | Molecular weight | Isoelectric point | Hydrophobicity (GRAVY)* | Dominant amino acids |
| --- | --- | --- | --- | --- | --- | --- |
| *Anopheles culicifacies* | 216 | - | 24.6 kDa | 5.10 | -0.395 | Ala (A), Glu (E), Leu (L)/ Gln (Q) |
| *Anopheles stephensi* | 215 | 94.44 % | 24.6 kDa | 5.19 | -0.383 | Ala (A), Leu (L), Glu (E) |
| *Anopheles gambiae* | 216 | 91.67 % | 24.6 kDa | 5.07 | -0.366 | Ala (A), Leu (L), Gln (Q) |
| *Homo sapiens* | 183 | 31.38 % | 21.2 kDa | 5.31 | -0.773 | Leu (L), Glu (E), Asp (D) |
| *Rhipicephalus sanguineus* | 172 | 36.96 % | 19.9 kDa | 5.13 | -0.685 | Leu (L), Glu (E), Ala (A) |
| *Drosophila melanogaster* | 205 | 42.65 % | 23.1 kDa | 5.59 | -0.240 | Leu (L), Ala (A), Glu (E)/Lys (K) |
| *Escherichia coli* | 158 | 22.73 % | 18.5 kDa | 4.79 | -0.472 | Leu (L), Glu (E), Asp (D) |
| *Arabidopsis thaliana* | 259 | 26.39 % | 29 kDa | 6.16 | -0.269 | Leu (L)/Ser (S), Glu (E), Lys (K)/ Val (V) |

\*Negative values indicate hydrophilic property

In a recent noteworthy study on ticks, ferritin knockdown has been shown to be lethal, underscoring its indispensable role in blood-feeding species^14–16^. Given the hematophagous nature of mosquitoes, a similar essential role is predicted for ferritin in their survival, reproduction, development and iron management. In *Aedes aegypti*, ferritin has been implicated in both iron storage and transport functions that appear to be uniquely adapted within insect ferritins^20,21^. However, despite its physiological importance, *Anopheles* ferritin remains poorly characterized. Interestingly, recent studies in the ESCAPE group of bacteria have identified the ferritin–ferredoxin interface as a promising drug target ^22^, further highlighting the potential of ferritin and its interactions with other proteins as a molecular target in mosquitoes for vector control strategies.

In this study, we implemented a multidisciplinary approach to investigate ferritin and the broader iron metabolic pathway in *Anopheles* mosquitoes. Given that the heavy-chain subunit plays a pivotal role in iron oxidation and storage, and is highly conserved across *Anopheles* species, we focused our efforts primarily on this subunit to explore its structural and functional uniqueness in comparison to bacterial and human ferritins using diverse computational and experimental approaches. Through comprehensive phylogenetic and motif analyses, we uncovered the chimeric nature of mosquito ferritin heavy-chain homologs, which retain conserved motifs from both major ferritin subfamilies, classical ferritins and bacterioferritin’s ^23^. Remarkably, we identified putative haem-binding residues in *Anopheles* ferritin, a feature characteristic of bacterioferritin’s but lost during ferritin evolution in higher eukaryotes. Consistent with this observation, our binding assays revealed that *Anopheles culicifacies* ferritin exhibits even higher affinity for hemin than bacterial ferritin, highlighting a unique evolutionary adaptation.

To experimentally validate our other computational predictions, we recombinantly expressed and purified *Anopheles culicifacies* ferritin, confirming that it forms a stable 24-mer cage structure with ferroxidase and hemin binding activity. We further generated specific antibodies against the heavy-chain subunit to examine its localization and expression dynamics during blood feeding. Using immunological assays, including immunofluorescence (IFA) and ELISA, we established the spatiotemporal expression profile of ferritin in female *Anopheles* mosquitoes post-blood meal (PBM). Iron deprivation experiments further demonstrated a strong correlation between ferritin expression and iron availability, revealing that disruption of iron homeostasis significantly impairs oocyte development and reproductive success.

Collectively, our findings establish ferritin as an evolutionarily distinct and functionally indispensable protein in mosquito iron metabolism and reproductive biology.As per existing reports, this is the first study to investigate the critical role of iron metabolism in both mosquitoes and *Plasmodium* during the mosquito stage, while evaluating the potential of iron chelators as transmission-blocking agents.

## Results

### Phylogenetic Divergence of *Ac*FtnHCH with Conserved Iron-Binding Residues

As previously mentioned, insect ferritins remain largely unexplored. To date, only one insect ferritin has been characterized in detail, revealing a unique architecture composed of an equal number of heavy (H) and light (L) chain subunits (12 each) forming the protein cage ^18^. Given the remarkable diversity of insects and the fact that *Anopheles* mosquitoes alone comprise more than 500 species we initiated our study by examining the evolutionary relevance of ferritins, with a particular focus on mosquitoes ^24^. To trace the evolutionary divergence of *Ac*FtnHCH, we performed a comprehensive phylogenetic analysis using sequences homologous to the human ferritin heavy chain, collected across diverse species, orders, and classes, with emphasis on insects and mosquitoes. The resulting phylogenetic analysis revealed a high degree of sequence conservation among members of the two Culicidae subfamilies Anophelinae (*Anopheles*) and Culicinae (*Aedes*, *Culex*) (**Figure S1A & B**). When we extended the comparison to a broader phylogenetic context that included representatives from various insect orders, we found that members of the family Culicidae (*Anopheles* and *Aedes*) formed a cohesive and distinct clade, clearly separated from other hematophagous insects and even from other dipterans such as *Drosophila melanogaster* and *Glossina morsitans* (also a blood-feeding species), as well as from the order Ixodida (class Arachnida) (**Figure S1C & D**). The observed divergence of ferritin sequences across hematophagous insects likely reflects lineage-specific ecological adaptations and host–parasite interactions. Notably, in Culicidae, only females are facultative blood-feeders required for egg maturation, whereas in other hematophagous taxa, both sexes are obligate blood-feeders ^11^. At a higher taxonomic level, Culicidae members consistently formed a distinct clade. Likewise, vertebrate and prokaryotic ferritins clustered separately, corresponding to two major ferritin subfamilies: classical ferritin and bacterioferritin ^12,23,25^ (**Figure 1A**). Structurally, ferritin is a cage-like protein exhibiting 4–3–2 symmetry and composed of 24 subunits (monomers). Each ferritin monomer typically adopts a characteristic four-plus-one (4 + 1) α-helical arrangement, with distinct conserved motifs that define individual subfamilies. Interestingly, mosquito ferritin exhibits feature characteristic of both bacterial and classical ferritins, indicating a chimeric nature. Specifically, it contains the E-6-Y (residues 58–65), EE-2-H (residues 92–96), and E-6-L/I (residues 147–154) motifs located in helices A, B, and C, respectively (**Figure 1B**), consistent with the classical ferritin heavy chain ^23^. In addition, it also possesses the E-2-H (residue no. 180-183) motif in helix D, typically seen in bacterioferritin ^23^ (**Figure 1A**). These features collectively position mosquito ferritin as a potential evolutionary intermediate between the two ferritin subfamilies. Since these conserved motifs are directly or indirectly involved in metal binding (i.e., the formation of ferroxidase sites), we superimposed the modelled *Ac*FtnHCH structure with several Fe²□-bound ferritin structures available in the Protein Data Bank (PDB) structure (Hftn-PDB ID: 4Y08, Bftn PDB ID: 1NF6). Following superimposition, we analysed the residues interacting with iron ions. Typically, ferritins possess two metal-binding sites enriched in glutamate (E) and histidine (H) residues for coordination. The monomer of *Ac*FtnHCH (∼26 kDa) showed a closer structural alignment with human H-chain ferritin (∼21 kDa) (RMSD- 0.56Å) than with bacterioferritin (∼18 kDa), (**Figure S2A**). Although the superimposition of ferritin monomers from diverse species including human, chicken, frog, moth, and mosquito revealed a considerable structural similarity and conservation across taxa (RMSD of <0.6Å) (**Figure S2B**).

**Figure 1.**
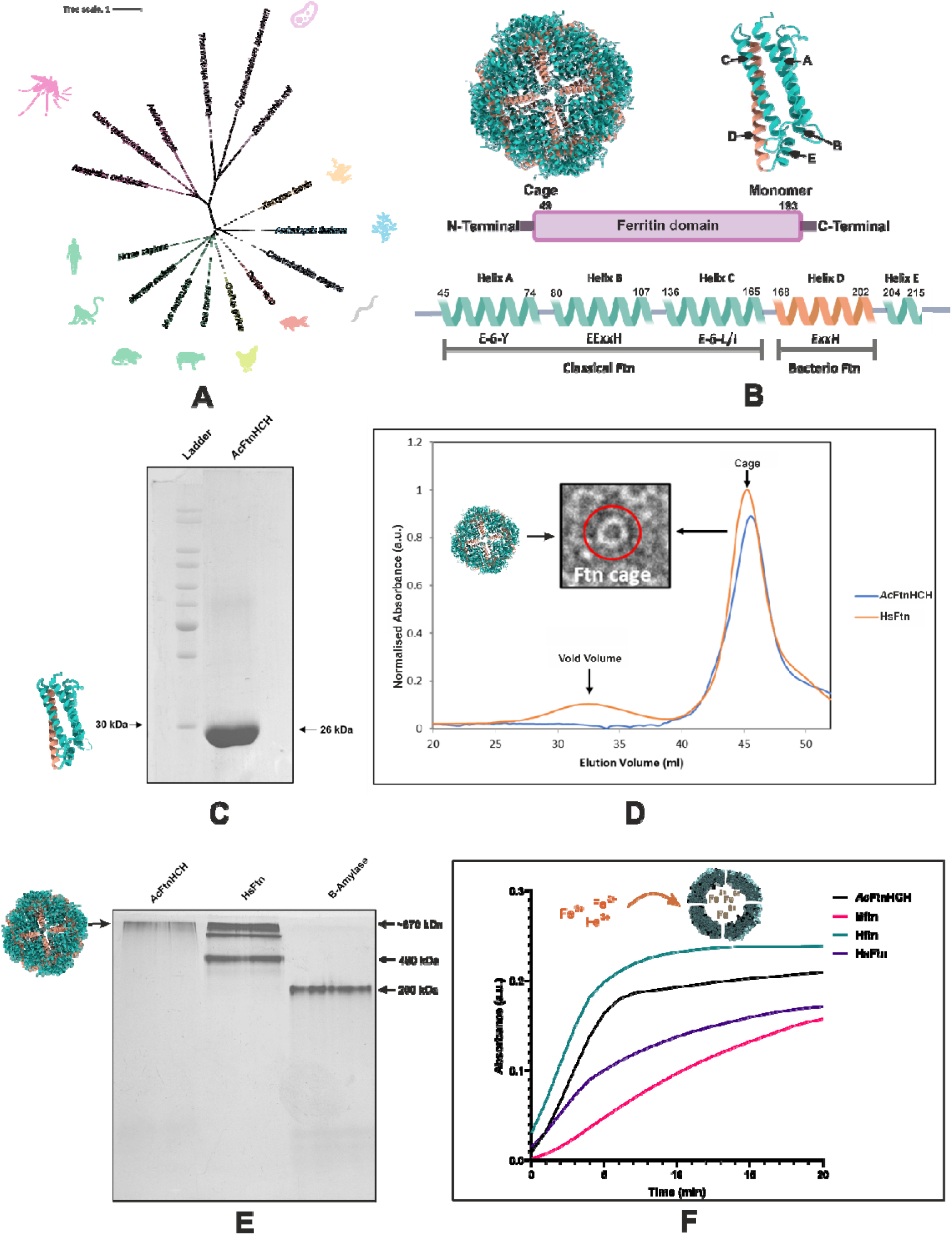
Sequence, structural, and evolutionary analysis of *Ac*FtnHCH and other ferritin family proteins. (A) Maximum likelihood phylogenetic tree (Le Gascuel substitution model, 1000 bootstrap replicates) depicting evolutionary relationships of ferritin proteins across diverse taxa. (B) Top left: Cage structure of *Ac*FtnHCH showing 24 subunits. Top right: Monomeric form highlighting chains, helices, and the ferritin domain spanning N- to C-terminus (below), Bottom: Schematic of *Ac*FtnHCH monomer illustrating its chimeric features with classical ferritin and bacterioferritin motifs. (C) SDS-PAGE analysis of refolded *Ac*FtnHCH. (D) Size exclusion chromatography (SEC) profile of *Ac*FtnHCH, with horse spleen ferritin (HsFtn) as a control, confirming multimer formation. Inset: Transmission electron microscopy (TEM) image of *Ac*FtnHCH (∼12 nm), demonstrating ferritin-like nanocage assembly. (E) Native PAGE analysis of refolded *Ac*FtnHCH (Lane 1), HsFtn (Lane 2), and β-amylase (Lane 3), confirming the 24-mer oligomeric state of *Ac*FtnHCH. (F) Iron oxidation kinetics were monitored at 310 nm to compare the ferroxidase activity of *Ac*FtnHCH with human heavy chain ferritin (Hftn), horse spleen ferritin (HsFtn), and bacterioferritin (Bftn). The 310 nm peak reflects Fe³□ formation as Fe²□ is oxidized at the ferroxidase center of the ferritin heavy chain, indicating the rate and efficiency of iron oxidation among the tested ferritins.

The ferritin monomer contains a di-iron binding site, with conserved iron-coordinating residues identified across different ferritin types. In human H-chain ferritin (PDB: 4Y08), these residues include Glu28, Glu63, His66, Glu108, and Gln142; in *AcFtnHCH*, they include Glu58, Glu93, His96, and Gln182; and in bacterioferritin from *Desulfovibrio desulfuricans* (PDB: 1NF6), they include Glu23, Glu56, His59, Glu98, Glu132, Glu147, and His135 ^26^ (**Table S1**). Bacterioferritin, in addition to iron binding, is also known for its haem-binding capability a feature that distinguishes it from classical ferritins. An additional haem-binding residue, Met57, located within the E-2-H motif of the B helix, facilitates haem association at the dimeric interface between the A and B helices in the haem-binding subunits of bacterioferritin ^27^. Interestingly, this residue is replaced by Arginine in both human H-chain ferritin and *Ac*FtnHCH, resulting in minimal haem binding.

Notably, superimposition revealed that two methionine residues, Met66 and Met98, located near the E-6-Y and EE-2-H motifs in helices A and B of *Ac*FtnHCH, lie within ∼2 Å of the haem molecule in bacterioferritin, closely resembling the arrangement observed in bacterioferritin and suggesting a potential haem-binding interaction (**Figure S2C**). This interaction was further validated by MST binding studies using hemin, a haem analogue. Interestingly, *Ac*FtnHCH exhibited almost two-fold higher binding to hemin compared to bacterioferritin, with Kd values of 67.8 ± 6.5nM and 136.5 ± 21.9 nM for *Ac*FtnHCH and bacterioferritin, respectively (**Figure S2D**). All these findings suggest that, despite evolutionary divergence, the core iron-binding residues of ferritin are highly conserved. Although helix D (E-2-H) and haem-binding features in mosquito ferritin show some resemblance to bacterioferritin, its iron-coordinating residues align more closely with those of classical ferritin. Nevertheless, the distinctive motifs in mosquito ferritin needs further structural investigation to clarify their specific roles in iron-binding dynamics.

### Structural and Functional Characterization of Recombinant *Ac*FtnHCH

To characterize the protein at the molecular level, particularly its cage formation and stability and also to generate antibodies for physiological studies, recombinant *Ac*FtnHCH was expressed and purified in *E. coli*. The purified protein migrated as a band of ∼26 kDa on SDS-PAGE. Notably, substantial proportion of the protein was localized in the insoluble fraction, a phenomenon also reported for other insect ferritins also ^28^. Protein was purified from insoluble fraction and refolded (**Figure 1C**). Subsequently, size exclusion chromatography (SEC) of the refolded protein showed an elution profile similar to that of HsFtn, further confirming multimer formation (**Figure 1D**). The characteristic cage-like architecture of ferritin was visualized using transmission electron microscopy (TEM). When stained with uranyl acetate, *Ac*FtnHCH exhibited donut-shaped spherical structures with dark centres, indicative of the apo-form (**Figure 1D**). This appearance arises as uranyl acetate penetrates the central cavity of the ferritin cage, producing negative staining and a contrast-enhanced hollow core ^29^. The protein was also analysed by native PAGE alongside controls (HsFtn, horse spleen ferritin, and β-amylase). The expected size of the *Ac*FtnHCH monomer is ∼26 kDa, and as ferritin assembles into a 24-mer, the corresponding cage protein should exceed ∼624 kDa ^30^. This was consistent with the band pattern observed for refolded *Ac*FtnHCH in the native-PAGE (**Figure 1E**).

We also performed far-UV circular dichroism (CD) spectroscopy to validate the conformation of *Ac*FtnHCH, as the protein was purified from inclusion bodies, and also to assess its thermal stability. The CD spectrum exhibited negative peaks at 208 and 222 nm, indicating that the protein predominantly adopts an α-helical structure, consistent with the canonical 4 + 1 helical arrangement of ferritins ^25^ (**Figure S3A**). To evaluate thermal stability, the protein was exposed to a temperature ramp from 25□°C to 90□°C in 5□°C increments. No signs of denaturation were observed up to 70□°C, and although partial unfolding occurred at higher temperatures, the protein remained largely intact even at 90□°C, indicating proper folding and high thermostability (**Figure S3B**). To further confirm correct folding, tryptophan-based fluorescence spectroscopy was used. The refolded *Ac*FtnHCH displayed a λmax of 335 nm, consistent with a folded protein (typically 330–335 nm for folded versus 350–355 nm for denatured proteins). Thermal denaturation monitored via fluorescence produced a similar stability profile as observed in CD spectroscopy. Importantly, upon cooling back to 25□°C, ferritin exhibited self-refolding, with a slight red shift to a λmax of 344 nm, indicating that unfolding is reversible (**Figure S3C**). Similarly, to assess the effect of chemical denaturants on refolded *Ac*FtnHCH, we performed a guanidine hydrochloride (GuHCl) denaturation assay monitored by tryptophan fluorescence. The refolded *Ac*FtnHCH remained stable up to 1.5 M GuHCl. However, beginning at 2 M, a red shift in λmax was observed, indicating partial unfolding of the protein, with complete denaturation occurring at 5 M GuHCl (**Figure S3D**).

The heavy chain of ferritin always carries a ferroxidase centre responsible for catalysing the oxidation of Fe²□ to the more inert Fe³□ form. Since *in-vitro* refolding can sometimes compromise protein functionality, we assessed whether the refolded *Ac*FtnHCH retained its enzymatic activity by performing iron oxidation kinetics. For comparison, human H-chain ferritin (Hftn), horse spleen ferritin (HsFtn), and bacterioferritin (Bftn) were included as controls (**Figure 1F**). In each case, 2 μM protein samples were incubated with 200 μM FeSO□, and absorbance at 310 nm was monitored over 20 minutes at 1-minute intervals ^27^.

The Km value for Hftn was the lowest (2.386 ± 1.396 μM), followed by *Ac*FtnHCH (3.602 ± 0.215 μM) and HsFtn (5.987 ± 1.815 μM), which predominantly contains L-chain subunits. Bftn exhibited the highest Km (52.53 ± 13.088 μM) and required an increased FeSO□ concentration (500 μM) to achieve measurable activity. The catalytic efficiencies (Vmax/Km) for Bftn, HsFtn, *Ac*FtnHCH, and Hftn were 0.0115 ± 0.001 min□¹, 0.076 ± 0.016 min□¹, 0.146 ± 0.008 min□¹, and 0.275 ± 0.004 min□¹, respectively, indicating substantially lower catalytic efficiency in Bftn compared to the other ferritins. This reduced efficiency likely reflects its lower iron storage capacity (∼2,500 Fe³□/cage) relative to mammalian ferritins (∼4,500 Fe³□/cage) ^12,31^. Interestingly, *Ac*FtnHCH did not display Km values as low as Hftn despite being a homopolymer of the H-chain. Moreover, *Ac*FtnHCH showed visible degradation at FeSO□ concentrations above 200 μM and was completely degraded at 1 mM, whereas HsFtn remained stable even at 1 mM (**Figure S3E**). This reduced stability and iron oxidation efficiency may result from subtle structural or conformational differences that prevent complete replication of the H-ferritin ferroxidase site. Additionally, its bacterioferritin-like D-helix may allosterically regulate ferroxidase activity, a possibility that warrants further investigation.

We also compared inter-subunit interactions across different ferritin heavy chains along their three symmetry axes (dimeric, 3-fold, and 4-fold) and observed notable variations in interface properties even between *Anopheles* ferritins (**Figure S4)**. In particular, *Anopheles culicifacies* ferritin (*Ac*FtnHCH) exhibited larger predicted interface areas at the dimeric and 3-fold symmetry axes (2534 Å² and 1024 Å², respectively) and stronger binding free energies (Δ_i_G = –28 kcal/mol and –8.2 kcal/mol). These interactions were accompanied by extensive hydrogen bonding (N_HB_) and salt bridges (N_SB_) interactions (N_HB_ = 11, N_SB_ = 5; N_HB_ = 8, N_SB_ = 3), indicating greater structural stabilization compared to Bftn. In contrast, Hftn showed a slightly larger 4-fold interface area (506 Å²) and stronger corresponding binding free energy (Δ^i^G = –6.2 kcal/mol; N_HB_ = 7) (**Figure S5)**. Furthermore, electrostatic surface potential analyses of human, mosquito, and bacterial ferritins revealed striking differences in charge distribution around the 3-fold, 4-fold pores and also the B-pore, an asymmetric iron transport channel between three monomers (primary iron transport channel in heterodimeric bacterioferritin) (**Figure S5)**. The 3-fold pore of mosquito ferritin displayed a highly negative electrostatic potential, indicative of an enhanced affinity for Fe²□ ion entry into the cavity. In contrast, human ferritin and bacterioferritin exhibited more neutral or less negative potentials at corresponding regions, suggesting species-specific differences in iron transport mechanisms. Notably, since the 3-fold pore is also implicated in phosphate entry and superoxide dismutase (SOD)-like activity, our analysis suggests that mosquito ferritin is less likely to exhibit such SOD-like behaviour ^32^.

### Spatiotemporal Distribution of *Ac*FtnHCH in Mosquito Tissues

Previous transcriptome-based studies have shown that ferritin expression is markedly higher in adult mosquitoes compared to their aquatic stages, and its tissue distribution dynamically changes before and after a blood meal, with notably elevated levels in the midgut and ovaries ^20,33,34^. This regulation appears closely linked to the iron requirements and/or iron load of these tissues, particularly during digestion of the blood meal. To gain deeper insight into this regulation, we performed immunofluorescence assays (IFA) to investigate the temporal and spatial expression pattern of *Ac*FtnHCH from 0 to 72 hours post-blood meal (PBM), at 24-hour intervals, using a polyclonal antibody raised against *Ac*FtnHCH.

Our results revealed a distinct rise in *Ac*FtnHCH expression in the abdominal region, encompassing the midgut and ovaries, beginning at 24 hours PBM, peaking at 48 hours, and subsequently declining by 72 hours, corroborating earlier observations in *Aedes aegypti* ^20,33^ (**Figure 2A**). This pattern was corroborated in dissected ovary samples collected at corresponding time points and further validated by (Enzyme-Linked Immunosorbent Assay) ELISA (**Figure 2B & 2C**). Furthermore, analysis of transcriptomic data from VectorBase revealed that under sugar diet expression of *Ac*FtnHCH is higher in the ovaries than in the midgut, and similarly, its expression is higher in adults compared to larvae (**Figure 2D**). To visualize iron localization, ovaries were stained with potassium hexacyanoferrate, which specifically binds to Fe³□ and forms an insoluble ferric ferrocyanide complex (Prussian blue) (**Figure S6**). Since ferritin predominantly stores iron in the Fe³□ form, the observed light greenish-blue coloration of the ovaries with the most intense staining at 48 hours PBM correlated strongly with the ferritin expression patterns detected in the IFA and ELISA assays.

**Figure 2.**
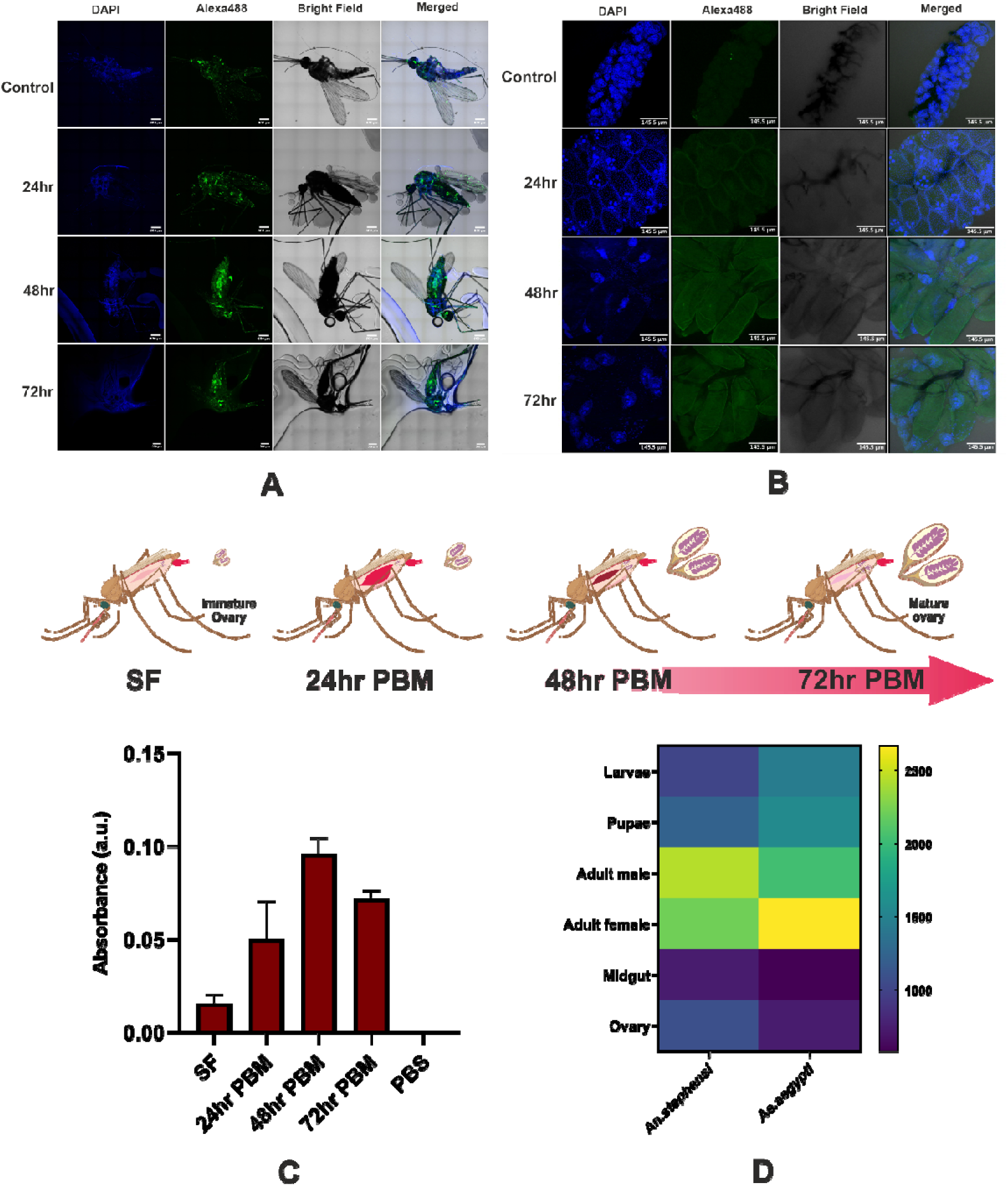
Expression and localization of *Ac*FtnHCH in *Anopheles culicifacies* females post blood feeding. (A) Whole-body immunofluorescence assay (IFA) showing *Ac*FtnHCH localization (green, Alexa488) and nuclei (blue, DAPI) in unfed control and at 24, 48, and 72 hours post blood meal (PBM). Fluorescence intensity increases after blood feeding. Scale bar: 500 μm.(B) Ovary IFA images depicting *Ac*FtnHCH distribution within ovarian tissue at the same time points. Ferritin expression increases over time. Scale bar: 145.5 μm; (C) ELISA quantification of *Ac*FtnHCH protein levels in sugar-fed (SF), PBS control, and at 24, 48, and 72 hr PBM, showing a time-dependent increase post blood feeding. Data are presented as mean ± SD from two independent replicates. (D) Comparative expression profile (TPM) of ferritin heavy chain homologues in *Anopheles stephensi* and *Ae. Aegypti* across different life stages and tissues, retrieved from VectorBase.

Together, these findings underscore ferritin’s pivotal role not only in iron storage but also in iron transport between the midgut and ovaries. The pronounced localization of ferritin in the ovary’s sites critical for oocyst formation and egg development highlights its physiological necessity in facilitating iron absorption and redistribution during periods of high post-blood meal iron influx.

### Iron Chelation Suppresses *Ac*FtnHCH Expression and Impairs Mosquito’s reproductive development

Ferritin plays a vital role in iron metabolism across diverse evolutionary lineages, serving as the primary iron storage and detoxification protein in most organisms ^30^. To validate the involvement of *Ac*FtnHCH in reproductive development of female mosquitoes, we conducted iron deprivation experiments using various iron chelators, including Deferoxamine, 2,2′-Bipyridyl, and β-Thujaplicin (**Table S2**). These chelators were tested at two distinct stages of the mosquito life cycle the larval (aquatic) and adult (terrestrial) stages, under both blood-fed and non-blood-fed conditions.

For larvicidal assays, ten third-instar larvae were kept in each well of a 24-well plate comprising 2 mL of Milli-Q water, with treatments performed in triplicate (**Figure 3A**). Increasing concentrations of the iron chelators were added, and larval mortality was recorded after 24 hours (**Figure 3B & C**). EDTA, a broad-spectrum metal chelator, was used as a control, while FeCl□ supplementation was employed to assess the effect of excess iron. Among the tested compounds, 2,2′-Bipyridyl exhibited the strongest larvicidal activity, with LD□□ values of approximately 91.8 μM for *Anopheles culicifacies* and 82.5 μM for *Anopheles stephensi* larvae. β-Thujaplicin showed LD□□ values of 260.4 μM for *Anopheles culicifacies* and >500 μM for *Anopheles stephensi*, while Deferoxamine exhibited LD□□ values of 371.8 μM for *Anopheles culicifacies* and >500 μM for *Anopheles stephensi*, indicating relatively low toxicity. In contrast, EDTA-treated larvae showed no mortality. Interestingly, larvae exposed to FeCl□ displayed LD□□ values of 121.3 μM for *Anopheles culicifacies* and 397.7 μM for *Anopheles stephensi*, indicating higher mortality compared to those treated with Deferoxamine or β-Thujaplicin, possibly due to increase in reactive oxygen species (ROS) levels generated by Fe³□. Additional compounds, including chrysin and gallium nitrate (Ga³□, a metal ion that competes with Fe³□ due to similar ionic radius and charge) (**Figure S7A**), were also tested but showed no significant larvicidal effect. Collectively, these findings indicate that the observed larvicidal activity is chelator-specific and primarily driven by the perturbation of intracellular iron homeostasis. Moreover, the lack of significant physiological effects from Ga³□ substitution suggests that Fe³□ plays a finely regulated yet nonredundant role in mosquito physiology.

**Figure 3.**
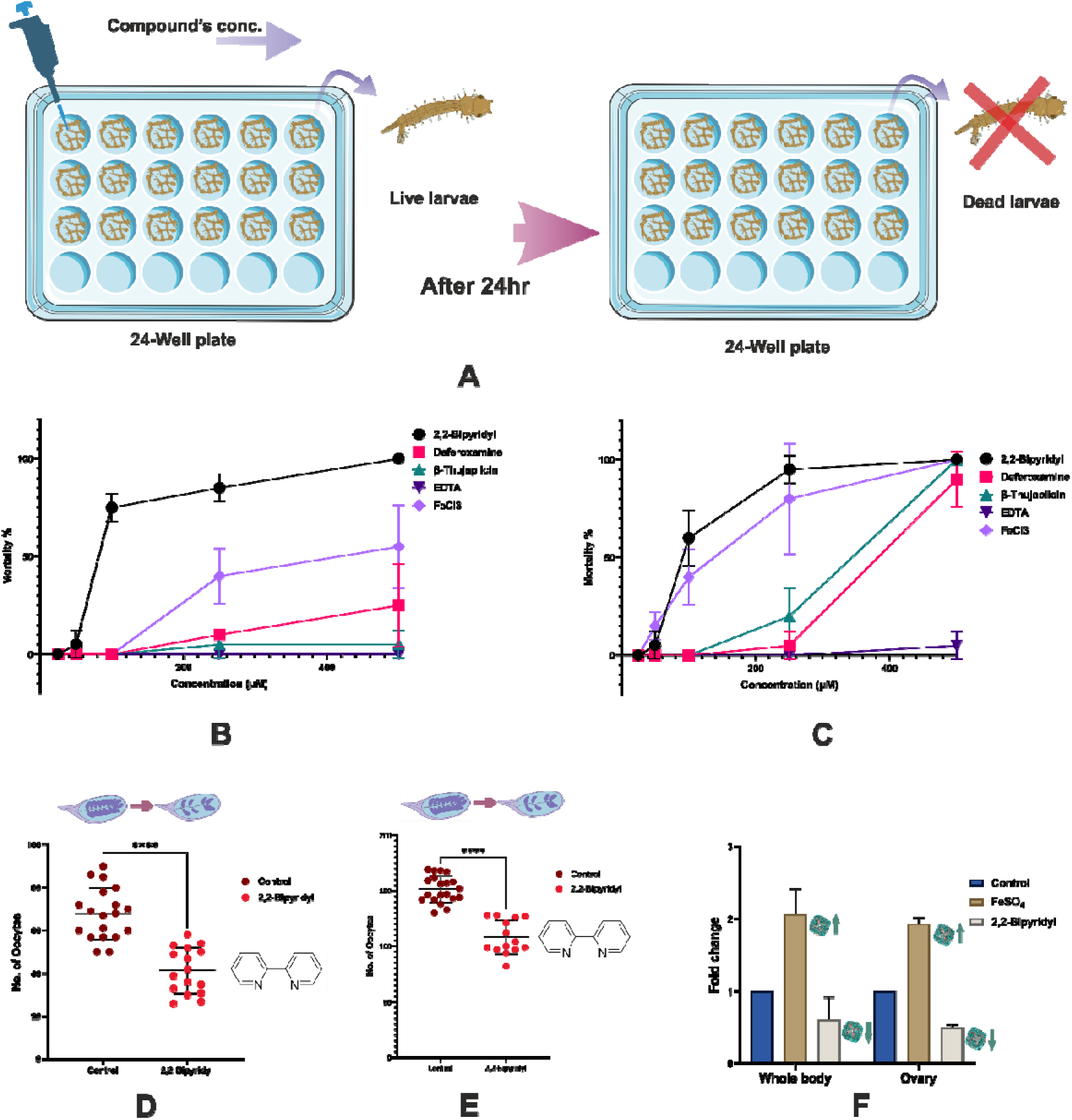
Larvicidal and ovicidal effects of different iron chelators in two different *Anopheles* mosquitoes; (A) Schematic representation of the larvicidal assay performed in the 24 well plate; (B C) Dose-dependent larval mortality (%) in *Anopheles stephensi* (B) and *Anopheles culicifacies* (C) following exposure to different iron chelators (2,2-Bipyridyl, Deferoxamine, β-Thujaplicin) as well as , EDTA, and FeCl_3_, showing highest efficacy for 2,2-Bipyridyl (n=30); (D, E) Dot plots showing significant reduction in the number of oocytes (D) *Anopheles stephensi*) and (E, *Anopheles culicifacies*) per female after 2,2-Bipyridyl treatment compared to control (****p < 0.0001, Mann–Whitney U test), with each dot representing a single female (n=20); (F) Bar graph depicting relative fold change in gene expression in whole body and ovary tissues of *Anopheles stephensi* following treatment with FeSO_4_ or 2,2-Bipyridyl compared to control, with data presented as mean ± SD levels of two independent replicates. infection, stained to visualize *Plasmodium* oocysts (indicated by circles and arrows). Oocyst numbers are visibly reduced in the 2,2′-Bipyridyl and β-Thujaplicin groups compared to controls.

Since female mosquitoes rely on host blood to obtain essential nutrients for egg development ^10^, and given that blood represents a major source of iron, we next investigated the effect of iron chelation on reproductive physiology. To this end, 3–4-day-old female mosquitoes were provided with 500 μM of various iron chelators via a cotton pad soaked in 10% sucrose solution, administered two days prior to and for three days following a blood meal. As anticipated, depletion of bioavailable iron resulted in a marked reduction in oocyte numbers, with 2,2′-Bipyridyl exhibiting the most pronounced effect reducing oocyte (immature eggs) counts by approximately -26.47 and -47.50 per mosquito in *Anopheles stephensi* and *Anopheles culicifacies*, respectively consistent with the larvicidal assay results (**Figure 3D & E; S7B)**. β-Thujaplicin exposure also resulted in the reduction of oocyte numbers, though the effect was comparatively moderate (−17 oocytes/mosquito in *Anopheles*. *stephensi*) (**Figure S7C)**. Analysis of differential gene expression further revealed that *Ac*FtnHCH was significantly downregulated in both whole-body and ovary samples of 2,2 Bipyridyl treated mosquitoes, likely reflecting mobilization of stored iron from ferritin under iron-deficient conditions. In contrast, *Ac*FtnHCH expression was upregulated in FeSO□-supplemented groups (**Figure 3F**). Together, these results establish a clear link between iron availability, ferritin expression, and reproductive success in mosquitoes. They underscore the central role of ferritin-mediated iron homeostasis in supporting oocyte maturation and overall reproductive fitness, highlighting iron metabolism as a promising target for vector control strategies.

## Discussion

Iron homeostasis represents a vital metabolic pathway in mosquitoes, as in other hematophagous vectors, ensuring proper iron utilization and detoxification. Ferritin, the principal iron-storage protein, has a major role in maintaining iron balance across species, and its evolutionary conservation underscores its physiological importance. Despite this, structural information on insect ferritin remains scarce currently limited to the ferritin structure from *Trichoplusia ni*. This structure revealed a tetrahedral cage symmetry ^18^, a feature shared with phytoferritin ^30^ but absent in mammalian ferritins. Interestingly, *Trichoplusia. ni* ferritin comprises an equal 12:12 ratio of H and L subunits, whereas ferritin in hematophagous insects such as mosquitoes may exist as homopolymers composed solely of either H or L chains. Generally, the H:L chain ratio is known to correlate with tissue-specific iron load, as demonstrated in mammals: ferritin in the heart, kidney, and brain is enriched in H-chains to enable rapid iron uptake and detoxification, whereas in the liver and spleen organs serving as iron reservoirs it is L-chain-rich, facilitating storage and slow release ^35^. Structural analyses of ferritins across species have further revealed notable differences in inter-subunit interactions along the dimeric, threefold, and fourfold symmetry axes (**Figure S4**). Even within a single genus, ferritins vary in interface area, bonding pattern, and binding energy, reflecting evolutionary fine-tuning of their structural stability and function.

Electrostatic surface analyses also indicate that the threefold axis pore in ferritin of *Anopheles* mosquito is markedly more negatively charged than those of bacterioferritin or human ferritin, suggesting an enhanced capacity for iron influx (**Figure S5A & B**). Because ferritin internalizes Fe²□ ions primarily through its threefold channels, the highly negative electrostatic potential at these sites in mosquito ferritin may facilitate efficient iron uptake, enabling adaptation to the elevated iron burden imposed by a haem-rich blood diet. In contrast to bacterioferritin heterodimers, where Fe²□ transport predominantly occurs through the B-pore ^36^, mosquito ferritin appears to utilize the threefold pore as its principal iron entry pathway. Interestingly, despite this difference, mosquito ferritin retains the ability to bind a haem moiety, similar to bacterioferritin. Such structural distinctions are likely to modulate ferritin’s overall stability, Fe²□ binding affinity, and ferroxidase activity, reflecting evolutionary optimization for iron handling in hematophagous insects.

Moreover, the chimeric nature of mosquito ferritin and the presence of potential haem-binding residues near the ferroxidase centre analogous to those facilitating electron relay and iron release in bacterioferritin further emphasize its unique structural and functional architecture ^37^. Collectively, these features suggest that mosquito ferritin has evolved specialized structural and electrostatic adaptations to efficiently regulate iron metabolism under the iron-rich conditions imposed by blood feeding. Iron serves as a high-energy nutrient that is indispensable for mosquito reproduction, being extensively utilized only by females during oogenesis. As demonstrated in iron chelator studies discussed earlier, iron deprivation markedly reduces oocyte numbers, underscoring the essential role of iron in reproductive success. Knockdown studies in tick studies has shown that, silencing of Fer1 (*Ac*FtnHCH ortholog) caused a greater reduction in oviposition and hatching rates than Fer2 knockdown ^16^, and also altered the midgut microbiome in some tick species (**Table 1**). This reinforces the predominant iron-storage and regulatory function of *Ac*FtnHCH orthologs relative to other ferritins in insects. Parallel findings in Drosophila show that heavy-chain ferritin knockdown leads to severe developmental deformities, such as malformed wings, whereas silencing the light chain produces negligible effects ^38^. Comparable trends have been observed across multiple insect taxa, further highlighting the conserved and critical role of heavy-chain ferritins in insect development (**Table 1**). In addition to ferritin, knockdown of other iron-associated genes such as transferrin and iron regulatory protein (IRP) also impacts reproductive fitness, though to a lesser extent ^15,16^. This indicates that while ferritin is not the sole determinant of iron delivery to developing tissues, it performs an essential role in maintaining iron homeostasis, mitigating oxidative stress via iron sequestration, and coordinating with other iron-regulatory pathways to support reproductive and developmental processes.

With the accelerating emergence of resistance to both insecticides and anti-plasmodial drugs, the need to identify novel targets and therapeutic molecules has become critical. Targeting essential metabolic pathways has historically yielded effective interventions, as evidenced by several drugs that maintain efficacy with a delayed onset of resistance ^39^.

In summary, this study reports the first evidence of the chimeric nature of mosquito ferritin, highlighting its potential haem-binding activity through integrated phylogenetic, structural, and biophysical analyses. We also established a strong link between iron availability and ferritin expression. Using iron chelators and targeted gene knockdown, we show that modulating ferritin levels directly impacts these critical biological processes.

### Methods Mosquito rearing

*Anopheles culicifacies* and *Anopheles stephensi* were reared in the insect rearing facility of Indian Council of Medical Research-National Institute of Malaria Research (ICMR-NIMR). The temperature and humidity of the insectary were maintained to 27° ± 1 °C and 80 % respectively, with 12:12 hours light/dark cycle. A mix of fish food and dog biscuits in 1:1 ratio was fed to the larvae, while 10% sucrose solution was fed to the adult mosquitoes through a soaked cotton pad. Mosquito rearing and maintenance protocols were authorized by the ICMR–NIMR Ethics Committee.

### Expression and purification

The construct containing the *Ac*FtnHCH sequence (VecorBase: A0A182M0V3) in the pRSET-A vector was acquired from Invitrogen, the Hftn construct (pET-A Vector) was obtained from BIOMATIK, Canada and the Bftn construct (pET-A Vector) was kindly provided by Dr. Arnab Basu, RKMVERI, West Bengal, India. Recombinant *Ac*FtnHCH, Hftn, and Bftn were expressed in BL21(DE3) cells (NEB). The cells were grown for 3 hours at 37 °C in LB medium, and at an OD_600_ of ∼0.6, the protein expression was initiated using 0.5 mM isopropyl β-D-1-thiogalactopyranoside (IPTG). After an overnight incubation at 16 °C, the culture was centrifuged at 6000 rpm for 20 minutes at 4 °C and the bacterial cell pellets were collected. Cells expressing *Ac*FtnHCH were resuspended in 8 M urea, 20 mM Tris, 250 mM NaCl, pH 8.0, and lysed by sonication. The lysate was centrifuged at 13,000 rpm for 30 minutes at 4 °C, and the resulting supernatant was incubated with Ni-NTA resin (Qiagen) at room temperature for 1 hour on a gel rocker for binding. The protein was purified using an Econo-Column® Chromatography Column (Bio-Rad) via an imidazole gradient (10–500 mM). Cells expressing Hftn and Bftn were lysed in lysis buffer (50 mM Tris, 300 mM NaCl and 1 mM PMSF; pH 8.0) through sonication after 20 minutes of incubation with 0.3 mg/ml lysozyme at room temperature. The supernatant obtained after the lysate was centrifuged at 13,000 rpm for 30 minutes at 4 °C was loaded onto a HisTrap™ HP column and purified using the ÄKTAprime Plus system (Cytiva) via an imidazole gradient method.

### Refolding and SEC

The purified recombinant *Ac*FtnHCH was dialyzed twice overnight using dialysis tubing cellulose membrane (Merck) in refolding buffer (50 mM Tris, 100 mM NaCl, 5% glycerol and 0.5 mM DTT; pH 8.0) with constant stirring at 4 °C. The protein to refolding buffer ratio was 1:1000 and it was then consecutively dialyzed in the buffer with the same composition minus DTT. After dialysis, the protein was concentrated and subjected to HiLoad® 16/600 Superdex® 200pg (Cytiva) size exclusion chromatography (SEC) to determine the oligomer formation. Horse spleen ferritin was used as a reference for comparison. The SEC elutes were analysed using a 7% Native PAGE gel alongside the horse spleen ferritin.

### TEM Imaging

Transmission electron microscopy (TEM) was used to determine the cage formation of the refolded *Ac*FtnHCH. Using JEOL JEM-1230 microscope operated at an acceleration voltage of 80–100 kV. Briefly, 3.5 µL (.25–0.5 mg/mL) of the sample was placed onto glow-discharged copper-Formvar grids (FC400Cu100, EM Resolutions Ltd, UK). For staining, 3.5 µL of 2% uranyl acetate was added after 1-minute of incubation, followed by another 1-minute incubation, and the surplus stain was removed.

### Biophysical and Functional Analysis of *Ac*FtnHCH

The proper folding of refolded *Ac*FtnHCH was assessed using CD and fluorescence spectroscopy. For secondary structure analysis, 4□μM protein in 50□mM phosphate buffer (pH 8.0) was analysed in the far-UV range (190–260□nm) using a JASCO-815 spectropolarimeter equipped with a Peltier thermostat. The thermal stability of the same protein sample was evaluated by thermal ramping in 5□°C increments up to 90□°C, with a 5-minute interval at each temperature. Intrinsic tryptophan fluorescence was then measured under the same buffer/protein conditions using an excitation wavelength of 292nm on an Eclipse Cary Varian UV-Vis spectrofluorometer (Varian Inc., Palo Alto, CA, USA) equipped with a Peltier temperature controller. Thermal stability was again monitored as in the CD experiment, followed by cooling the sample back to 25□°C to observe potential self-refolding. Additionally, GuHCl-induced stability was assessed by incubating the protein with increasing concentrations of GuHCl (0–5.5□M) for 30 minutes, followed by fluorescence measurement to detect changes in the tryptophan microenvironment. To observe iron oxidation kinetics, 2□μM protein was mixed with 200□μM FeSO□ in 100□mM HEPES,100mM NaCl buffer (pH 6.5). The oxidation of Fe²□ to Fe³□ was monitored at 310□nm for 20 minutes at 1-minute intervals. Readings were taken immediately after the addition of FeSO□ using quartz cuvettes in a Cary Varian Bio100 UV-Vis spectrophotometer at 25□°C. Kinetic parameters, including Vmax and Km, were determined by fitting the data to the Michaelis–Menten model using GraphPad Prism 9 (La Jolla, CA, USA).

### Murine Antibody production and Immunodetection analysis

Polyclonal antibodies against *Ac*FtnHCH were raised in ∼6-week-old female BALB/c mice at ICMR-NIMR, New Delhi, following standard procedures ^40^ and with pre-approval from the institute’s animal ethics committee (IAEC/NIMR/2019-1/09). These antibodies were used to detect *Ac*FtnHCH in the lysates of *Anopheles culicifacies* mosquitoes. For this, ovary lysates collected from adult females at 48 hours post blood meal (PBM), along with whole-body lysates from adult males, adult sugar-fed females, and adult blood-fed females, were separated on a 12% SDS-PAGE gel. The resolved protein bands were then transferred onto a nitrocellulose membrane (Bio-Rad) for western blot analysis, and was detected using *Ac*FtnHCH antibodies. For indirect ELISA, whole-body lysates from adult females collected at different time intervals post blood meal were used, with sugar-fed females serving as controls. The experiments were conducted using two independent biological replicates.

### Immunofluorescence Assay

To localize the protein within female mosquitoes, ovaries were dissected at various time points post-blood meal (24, 48, and 72 hours), with ovaries from sugar-fed females serving as controls. Sample preparation for the ovaries followed the method described in ^20^. Female adults were also collected at the same time points as the ovaries (3 mosquitoes for each timepoint), and whole-body sample preparation was carried out according to the previously described protocol ^41^, with minor modifications. The anti-mouse Alexa Fluor™ 488 secondary antibody and ProLong™ Gold Antifade Mountant was purchased from Invitrogen. The ovary samples were visualized using Leica TCS SP8 (AOBS- Acousto Optical Beam Splitter based), while the whole-body samples were visualized in Nikon A1R MP Laser Scanning Confocal Microscope.

### Prussian blue staining

To detect ferric iron (Fe^3+^) in the ovarian tissues, samples were collected at various time points post-blood meal (24, 48, and 72 hours), with ovaries from sugar-fed females serving as controls. The sample preparation followed the method described in ^20^. A freshly prepared Prussian blue staining solution consisting of 2% Potassium hexacyanoferrate (Sigma) and 2% HCl in 1:1 ratio was used for staining the tissues for 1hour at room temperature. The tissues were visualized under Olympus BX63 Light Microscope.

### Iron chelator treatment

#### Larvicidal

The 3^rd^ Instar larvae were transferred to a 24 well plate (10 larvae per well in triplicates) in 2ml/well of Millipore distilled water. The larvae were kept undisturbed for 15 minutes for acclimatization, and then the compounds (Deferoxamine, β-Thujaplicin, 2,2-Bipyridyl, EDTA, FeCl_3,_ Chrysin and Gallium nitrate) were added to each well at different concentration (500 μM, 250 μM, 100 μM, 50 μM, 25 μM) (**Table S2**). To rule out the mortality occurring due to the solvent used for dissolving the compounds, the highest vol. of solvent used per compound was taken as a control. The mortality rate was then counted 24 hours post treatment. The experiments were conducted using two independent biological replicates.

#### Ovicidal

The female mosquitoes (3-4 days post emergence) were fed with the 2,2-Bipyridyl (500 μM) and β-Thujaplicin (500 μM) mixed in 10% sucrose solution through cotton pad, 2 days prior and 3 days after the bloodmeal, the sugar fed females were used as a control. The mosquitoes were then dissected 72 hours post blood meal and their oocytes were counted under a microscope. The data were analysed with GraphPad Prism 9 software (La Jolla, CA, USA) using Mann–Whitney U test. The experiments were conducted using two independent biological replicates.

### Reverse Transcription-PCR Expression Analysis

For the qPCR analysis, the female *Anopheles stephensi* mosquitoes (A0A182XWX0) were fed with 2,2-Bipyridyl (500 μM) and FeSO_4_ (1mM), with sugar-fed females as a control. Total RNA was extracted from the whole body and ovaries of female mosquitoes dissected at 72-hours post-blood meal, using RNAiso Plus (Takara) as per the manufacturer’s protocol. For cDNA library construction, the PrimeScript™ 1st Strand cDNA Synthesis Kit (Takara) was used. qPCR was conducted on a BioRad CFX96 Touch Real-Time PCR. The primer sequences were as follows, with S7 used for internal normalization (**Table S3**). The primers were ordered from Eurofins. The experiments were conducted using two independent biological replicates.

### Bioinformatics

The amino acid sequences for the orthologs of *Ac*FtnHCH in sixteen highly evolvable species ^42^ were sourced from VectorBase( https://vectorbase.org/vectorbase/app/ ), the sequence for different orders were collected from NCBI(https://www.ncbi.nlm.nih.gov/ ) and the ortholog sequences for the different classes were obtained from UniProt( https://www.uniprot.org/) (**Table S4**). An unrooted phylogenetic tree was constructed using the Maximum Likelihood method with LG model in MEGA X software, with a bootstrap value of 1,000. The domain structure was analysed with InterProScan( https://www.ebi.ac.uk/interpro/search/sequence/ ), and the 3D structures for *Ac*FtnHCH and *As*FtnHCH were predicted and refined using trRosetta(https://yanglab.qd.sdu.edu.cn/trRosetta/ ) and GalaxyWEB(https://galaxy.seoklab.org/ ).The PDB structures for Hftn(4Y08), Bftn(1NF6) and Tnftn(1Z6O) were obtained from protein bank (https://www.rcsb.org/ ) and were modelled using PyMOL. Multiple sequence alignment was performed with JALVIEW.The interface interaction information between two monomers of ferritin cage at different axis were obtained from PDBePISA server (https://www.ebi.ac.uk/pdbe/pisa/ ).

## Data availability

All data are available in the main text or the supporting information

## Supporting information

This article contains supporting information.

## Author contributions

C. T., R. N. D., S. S., K. C. P., S. S. and S. C. writing–review & editing; R. N. D., S. S., K. C. P., S. S. and S. C. validation; R. P., S. B., R. N. D., S. S., K. C. P., S. S. and S. C. supervision; R. N. D., S. S., K. C. P., S. S. and S. C. resources; S. S., K. C. P. and S. C. visualization; C. T., R. P., S. B., B. G., A. K., A. G., and A. L. methodology; C. T., R. P., S. B., B. G., A. G., A. L. and A. K., investigation; C. T. and S. C. writing–original draft; C. T., R. P., and S. C. formal analysis; S.C. Funding acquisition; S. C. conceptualization.

## Notes

The authors declare no conflict of interest.

## Acknowledgments

We are grateful to BITS, Hyderabad, for providing the necessary research infrastructure. We also thank the ICMR–National Institute of Malaria Research for financial, academic, and research infrastructure support We acknowledge the Central Instrumentation Facility and University of Delhi for access to confocal microscopy facilities. We also thank Mr. Rajan Singh, Facility In-Charge, Confocal Microscope Facility, SNIoE, for his assistance with confocal imaging, which was performed in part at the DST-FIST Confocal Microscope Facility, SNIoE (Grant No. SR/FST/LS-1/2017/59(c)). We extend our sincere thanks to Dr. Arvind Sharma and Dr. Bhuvan Dixit for their technical guidance, and to Mr. Uttam Singh, Mr. Satya, and Mr. V. P. Singh for their assistance in insectary maintenance. We also thank Dr. Arnab Basu (RKMVERI) for providing the Bftn plasmid.

