## Supplementary material for "Mosquito Ferritin: A Haem-Binding Iron Store Required for Egg Development and Targetable for Malaria Control": bioRXIV SI.docx

**Index**

**1. Supporting information figures**

Figure S1. Phylogenetic analysis of mosquito ferritin heavy chain homologs------------------------------- 3

Figure S2. Structural analysis of mosquito ferritin heavy chain homologs------------------------------------ 4

Figure S3. Structural, thermal, and chemical stability analysis of refolded *Ac*FtnHCH.-------------------- 5

Figure S4. Comparative analysis of ferritin subunit interactions across symmetry axes.------------------- 6

Figure S5. Comparative analysis of the surface electrostatic potential of ferritin fold-axis pores

from different species..---------------------------------------------------------------------------------- 7

Figure S6. Prussian blue staining reveals temporal accumulation of iron in *Anopheles stephensi*

ovaries following a blood meal----------------------------**------------------**-------------------------- 8

Figure S7. Effect of iron chelation on the development of *Anopheles stephensi* ---------------------------- 9

**2.Tables**

Table S1 List of conserved iron-binding residues in Human heavy chain ferritin, *Anopheles*

*culicifacies*’s ferritin heavy chain homolog and bacterioferritin from *Desulfovibrio*

*desulfuricans*-----------------------------------------------------------------------------------------------10

Table S2 Chemical and structural profiles of iron chelators used in the study-------------------------------11

Table S3 List of all the primers used in the study----------------------------------------------------------------12

Table S4 Amino acid sequence of ferritin’s synthetic construct used in the study---------------------------12

Table S5 List of all the species and their UniProt ID used to generate the phylogenetic tree--------------13

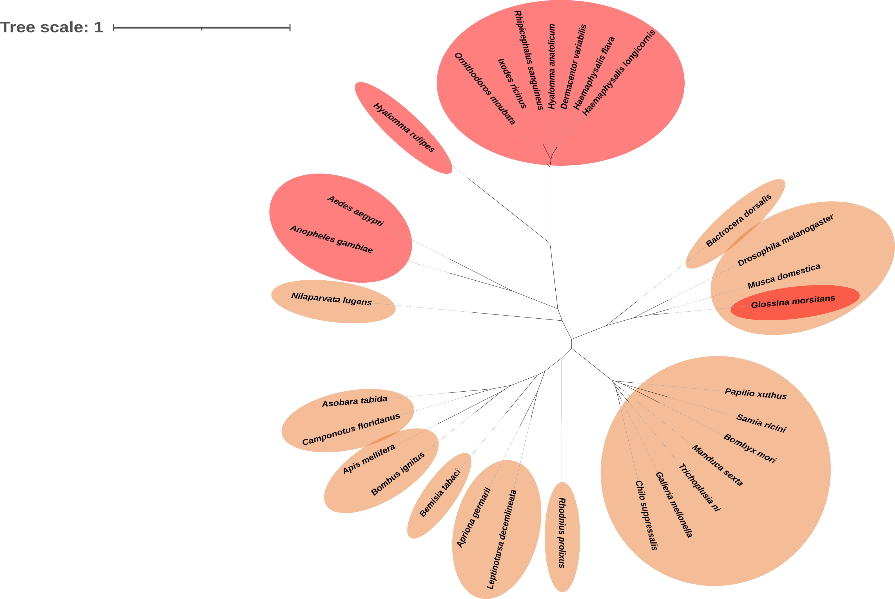

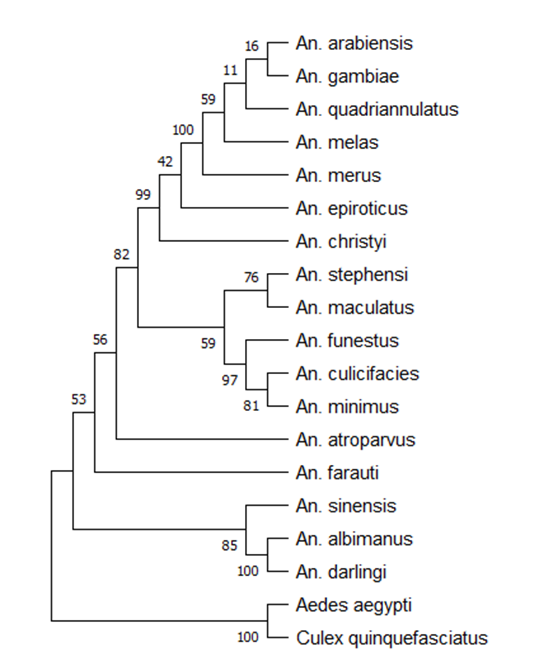

**C**

**A**

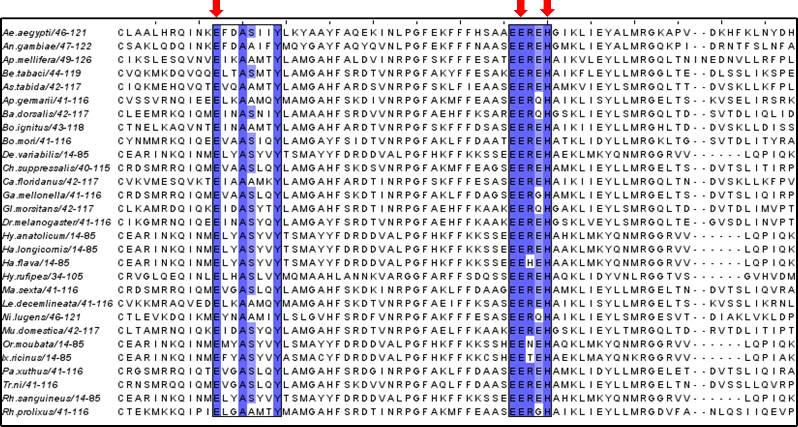

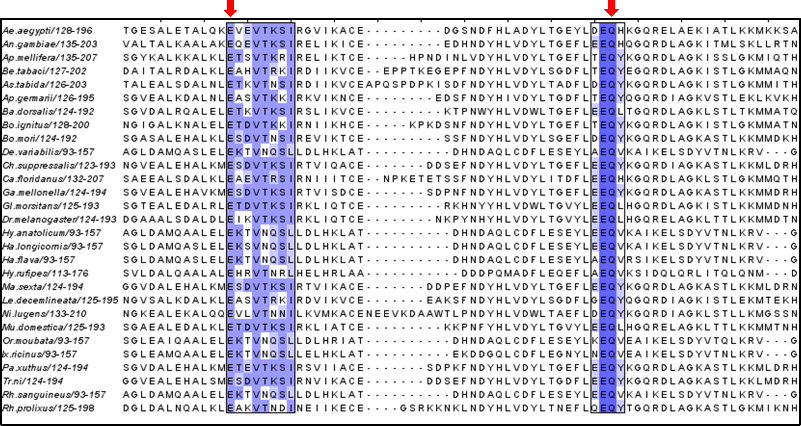

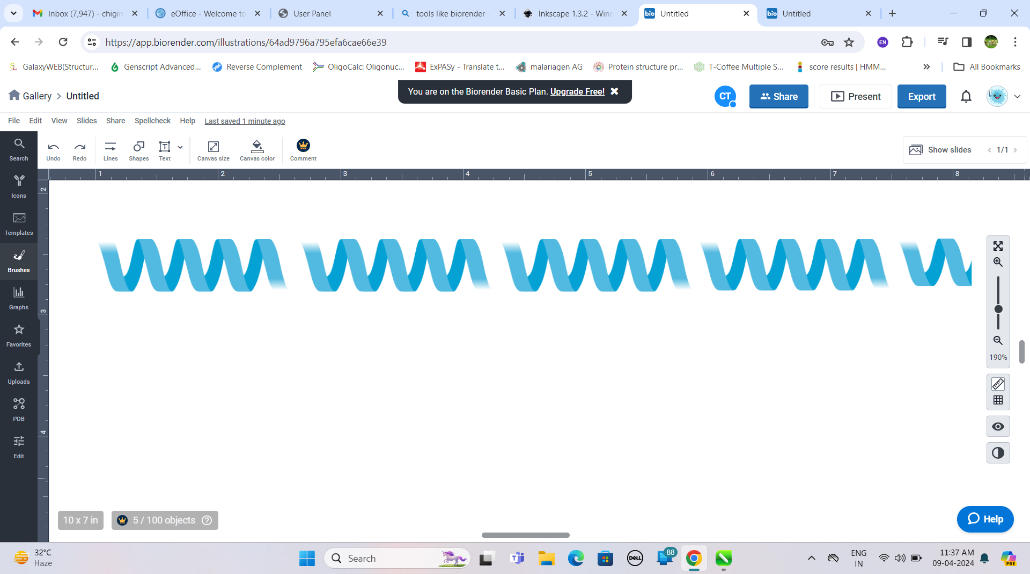

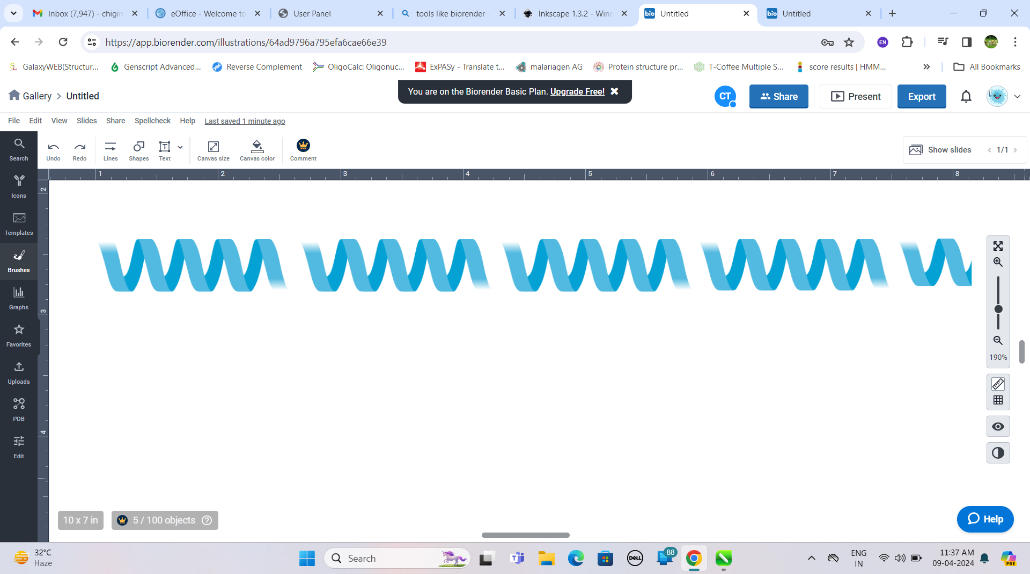

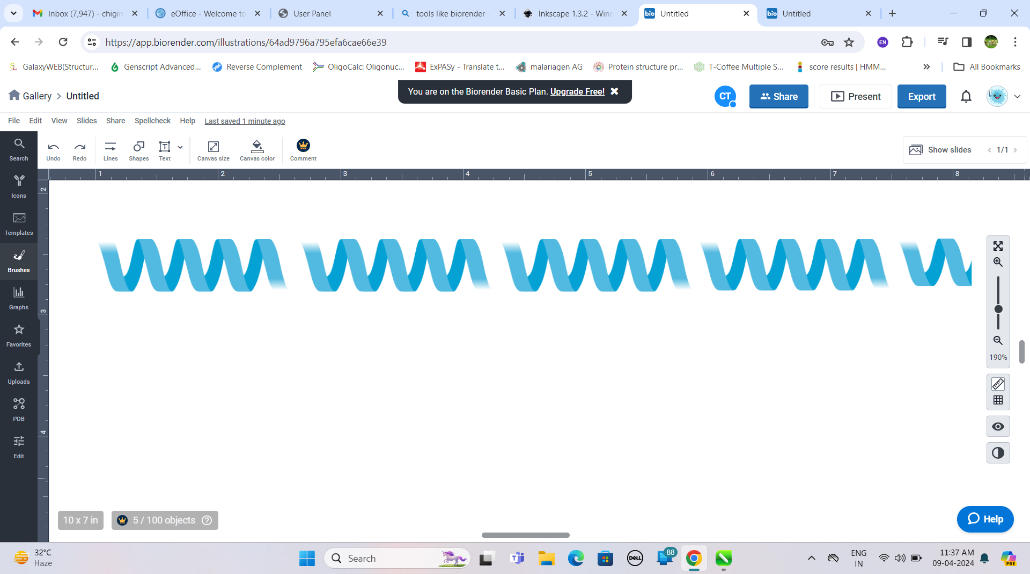

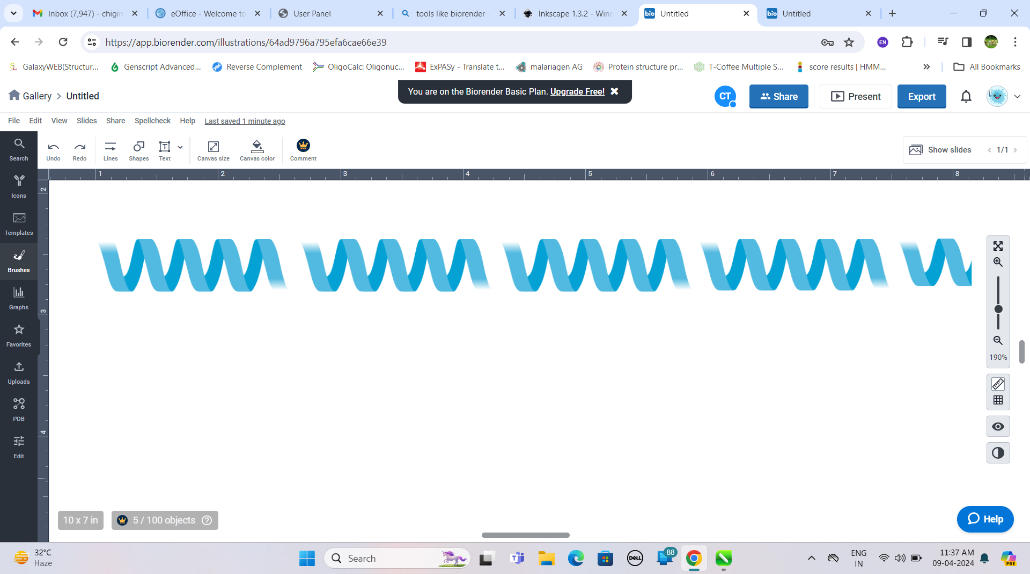

**Helix-D**

**Helix-C**

**Helix-B**

**Helix-A**

**B**

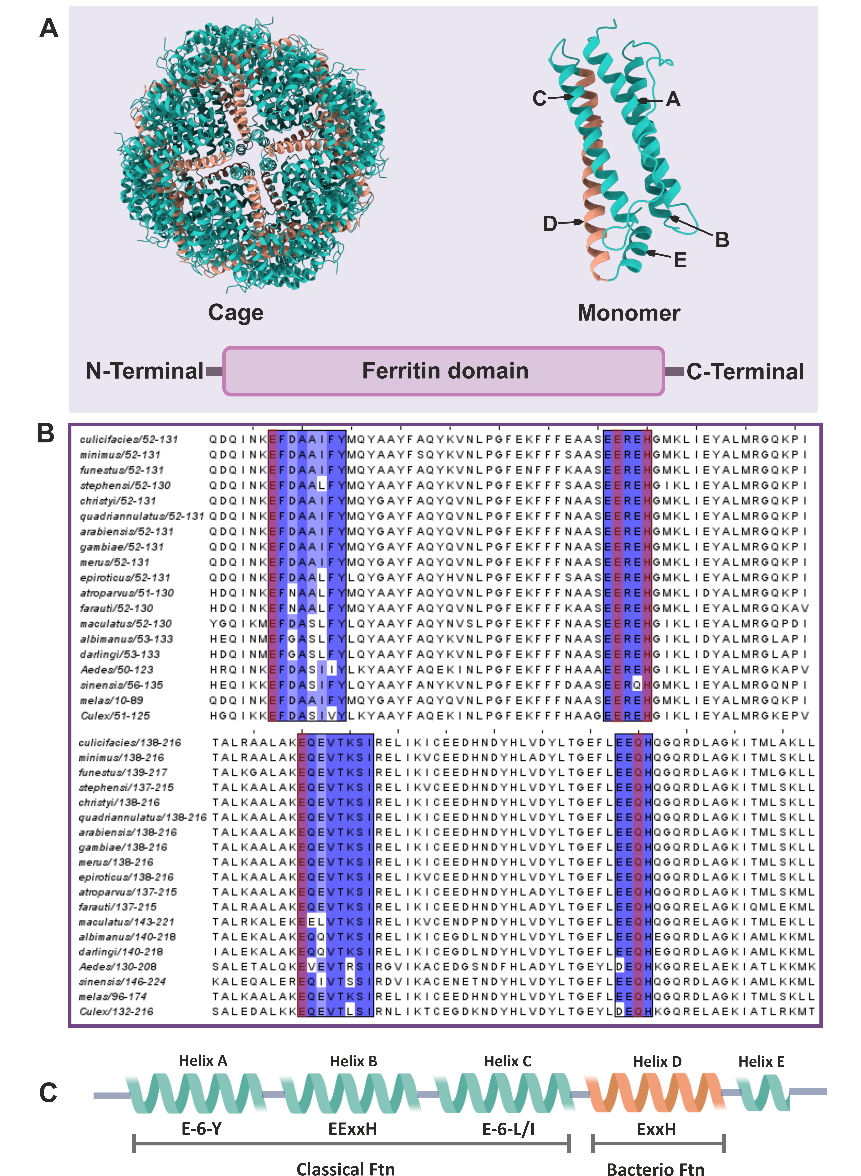

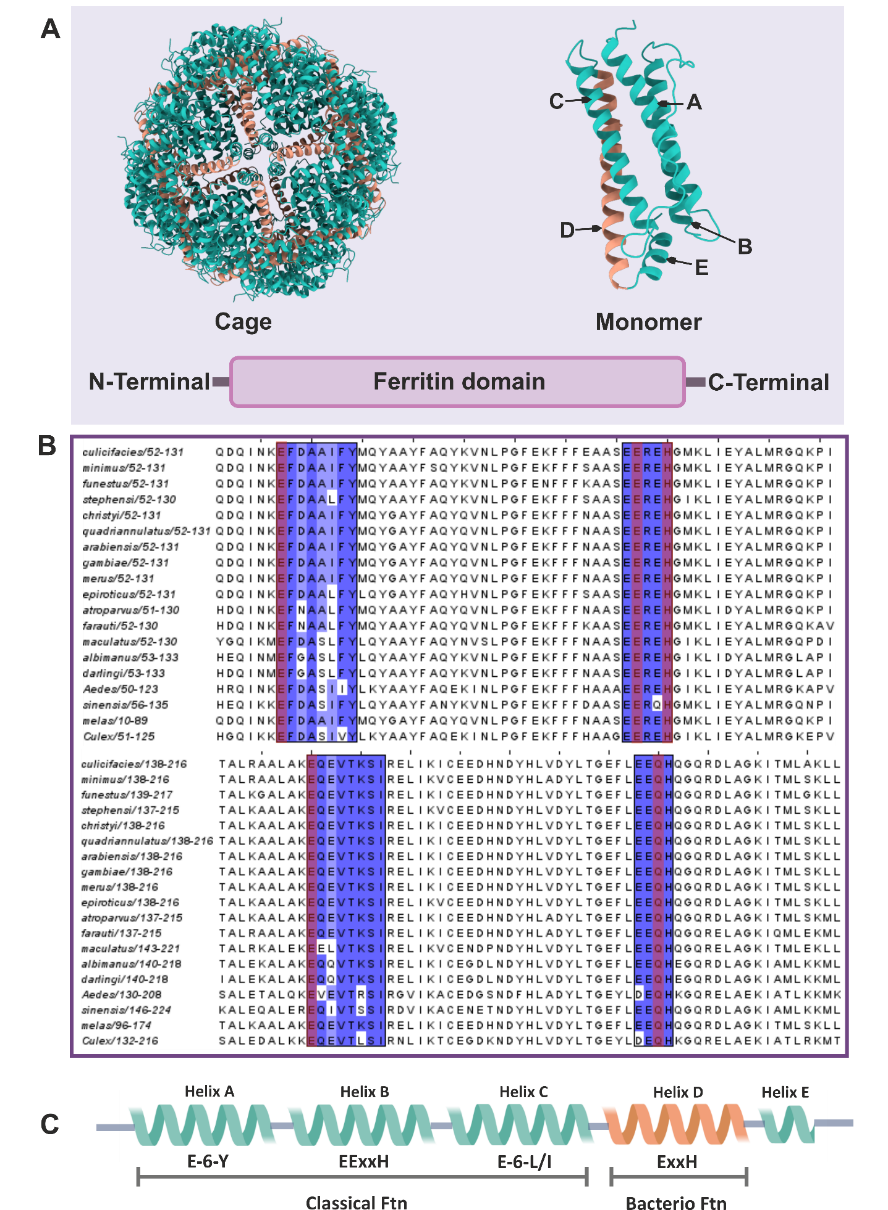

**D**

**Figure S1.** **Phylogenetic analysis of mosquito ferritin heavy chain homologs.**

(A) Maximum likelihood phylogenetic tree of ferritin heavy chain homologs from *Anopheles* and other mosquito species, illustrating evolutionary relationships within the genus and with other species (*Aedes aegypti* and *Culex quinquefasciatus*), bootstrap values are indicated at branch points. (B Multiple sequence alignment of the conserved motifs in ferritin heavy chain homologs from various Culicidae species, with darker blue indicating higher sequence conservation and red marking Fe²⁺-binding residues. (C Phylogenetic tree comparing insect ferritin heavy chain homologs sequences, demonstrating the distinct clustering of mosquito ferritins and their divergence from other insects as well as other blood-feeders (highlighted in red) (D) Multiple sequence alignment of ferritin heavy chain homologs from selected insect species, with conserved residues involved in iron binding (red arrow) and ferroxidase activity highlighted in blue.

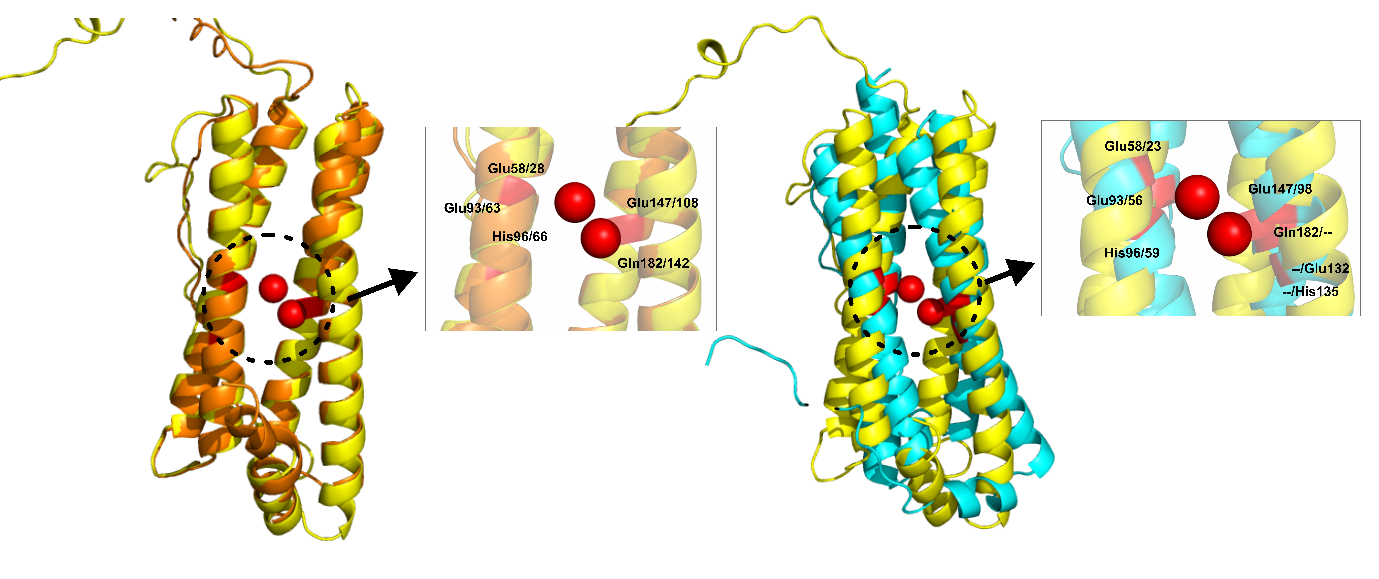

**A**

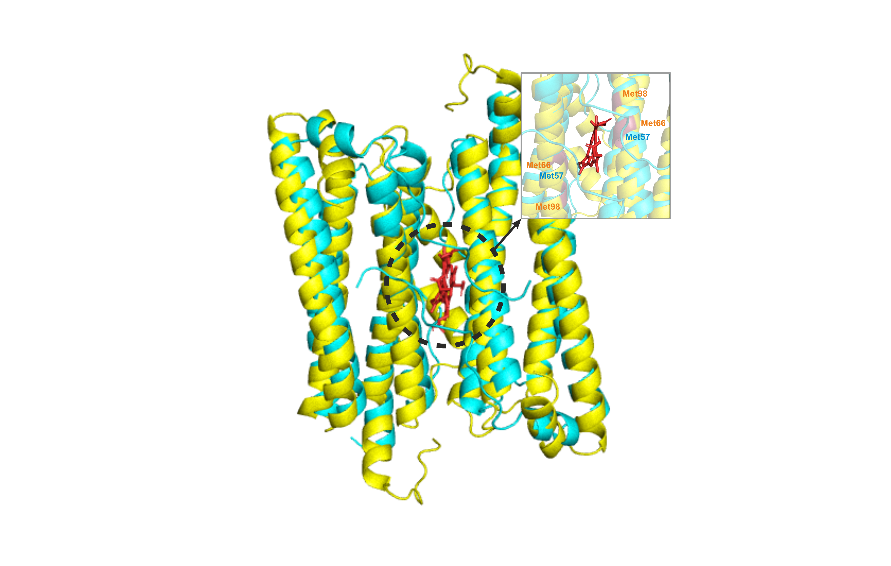

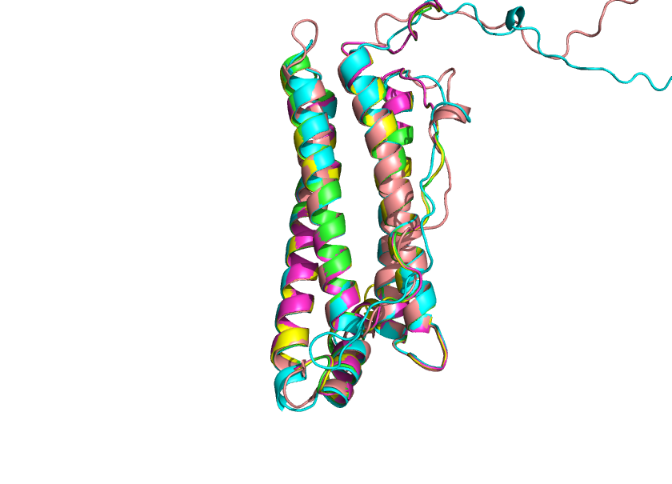

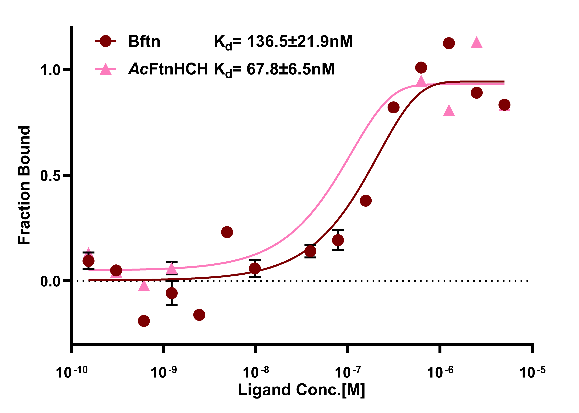

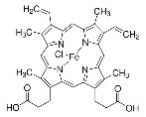

**B**

**D**

**C**

**Figure S2.** **Structural analysis of mosquito ferritin heavy chain homologs.**

(A) Structural superimposition of *Ac*FtnHCH with human H-chain ferritin (PDB: 4Y08) and bacterioferritin from *Desulfovibrio desulfuricans* (PDB:1NF4) in the presence of Fe²⁺ molecule, demonstrating conserved iron-binding residues among these proteins; (B) Structural superimposition of ferritin monomers from various species, including human, chicken, frog, moth, and mosquito, reveals a high degree of structural similarity across organisms. (C) Structural superposition of *Ac*FtnHCH (yellow) with bacterioferritin (cyan) reveals methionine residues positioned in close proximity to the haem molecule specifically, Met57 within the ferroxidase site motif of helix B in bacterioferritin, and Met66 and Met98 near the ferroxidase site motifs of helices A and B, respectively, in *Ac*FtnHCH, suggesting potential interactions with the haem group. (D) Microscale Thermophoresis (MST) binding curves show the determination of dissociation constants (Kd) for Bacterioferritin (K_d_ = 136.5 ± 21.9nM) and *Ac*FtnHCH (K_d_ = 67.8 ± 6.5nM) upon titration with hemin, indicating a higher binding affinity for *Ac*FtnHCH compared to Bacterioferritin.

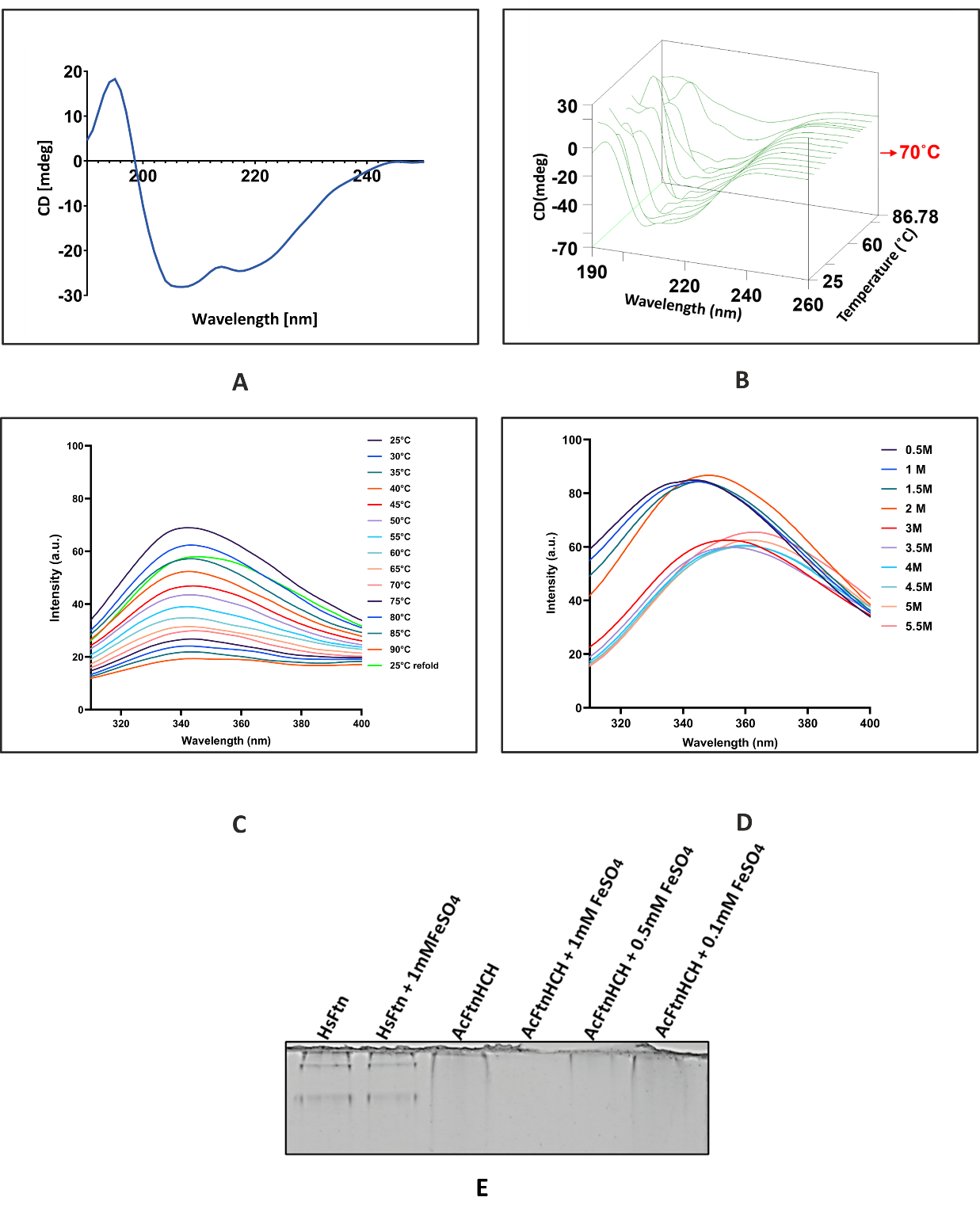

**Figure S3. Structural, thermal, and chemical stability analysis of refolded *Ac*FtnHCH.**

(A) Circular dichroism (CD) spectrum of refolded *Ac*FtnHCH, demonstrating a strong α-helical dominant secondary structure with characteristic minima near 208 nm and 222 nm; (B) CD thermal ramping of *Ac*FtnHCH from 25 °C to 90 °C with 5 °C increments, showing retention of helical structure up to ~70°C and indicating high thermal stability; (C) Tryptophan based fluorescence spectroscopy of *Ac*FtnHCH (contains single tryptophan) during thermal ramping, monitoring emission changes from 25 °C to 90 °C displaying high thermal stability and after lowering the temp back to 25°C ,the protein showed increase in fluorescence with a slight red shift in ʎ_max_ (max. fluorescence) suggesting partial reversible folding of the *Ac*FtnHCH; (D) Guanidine hydrochloride (GuHCl) denaturation test using tryptophan fluorescence, revealing the chemical stability of refolded *Ac*FtnHCH across increasing GuHCl concentrations; the refolded *Ac*FtnHCH showed stability upto 1.5M.However, starting at 2M, a red shift in ʎ_max_ was observed suggesting unfolding of the protein,with complete unfolding of the protein seen at 5M.(E) Native-PAGE analysis showing the stability of refolded *Ac*FtnHCH in the presence of increasing concentrations of FeSO₄ (0.1,0.5 and 1mM), along with Horse spleen ferritin (HsFtn) which used as control.

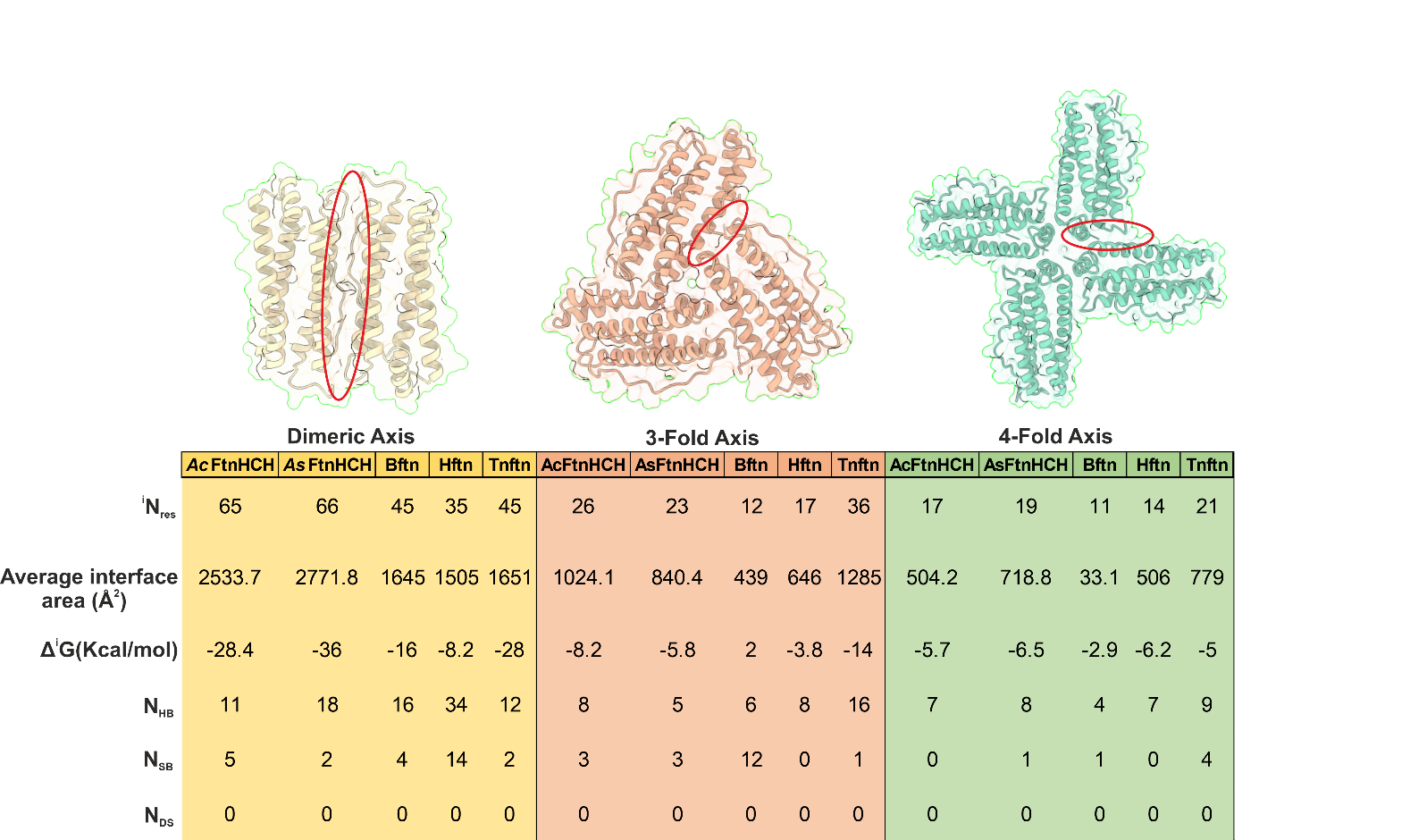

**Figure S4.** **Comparative analysis of ferritin subunit interactions across symmetry axes.**
Structural representation and interface characterization of ferritin homologues i.e., *Ac*FtnHCH (*Anopheles culicifacies*), *As*FtnHCH (*Anopheles stephensi*), Bftn (Bacterioferritin of *Desulfovibrio desulfuricans*), Hftn(Human), Tnftn(*Trichoplusia ni*) at the dimeric, 3-fold, and 4-fold symmetry axes. The table summarizes key parameters including the number of interacting residues (^i^N_res_), average interface area (Å²), binding free energy (Δ^i^G,Kcal/mol), hydrogen bonds (N_HB_), salt bridges (N_SB_), and disulfide bonds (N_DS_). Structural models above illustrate the subunit interfaces along each symmetry axis.The data were generated using PDBePISA.

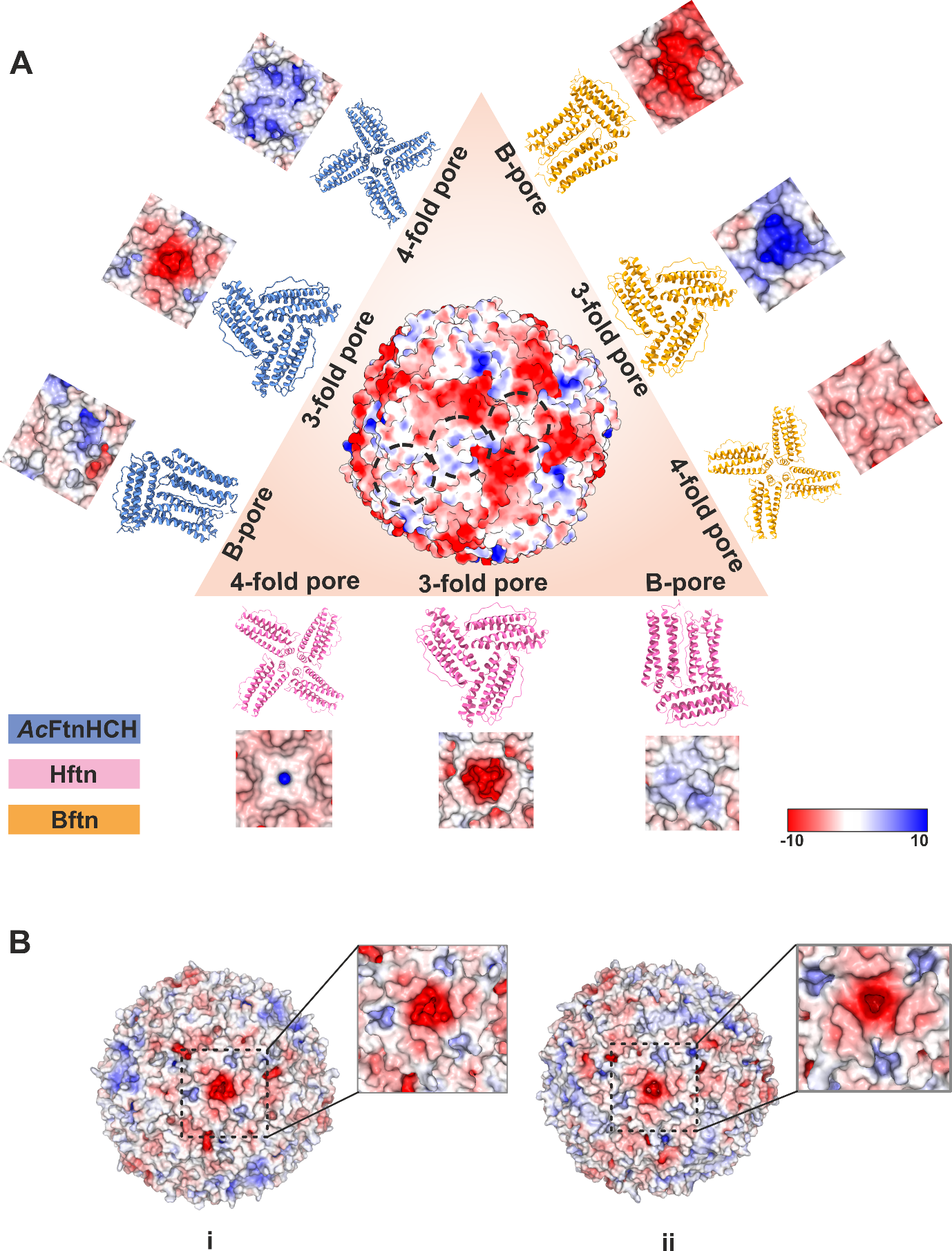

**Figure S5. Comparative analysis of the surface electrostatic potential of ferritin fold-axis pores from different species.** (A) Surface electrostatic potentials are shown for the ferritin heavy chain homolog from *Anopheles culicifacies* (*Ac*FtnHCH, blue), human (Hftn, pink), and bacterioferritin of *Desulfovibrio desulfuricans* (Bftn:PDB ID-1NF6, orange), highlighting the charge distribution on the pores at the B-fold, threefold, and fourfold axes. Red indicates negative potential (−10),white indicates neutral and blue indicates positive potential (+10) and was calculated using APBS module in Pymol; (B) Surface electrostatic potentials shown for the ferritin heavy chain homolog of i) *Anopheles culicifacies* and ii) *Anopheles stephensi*.

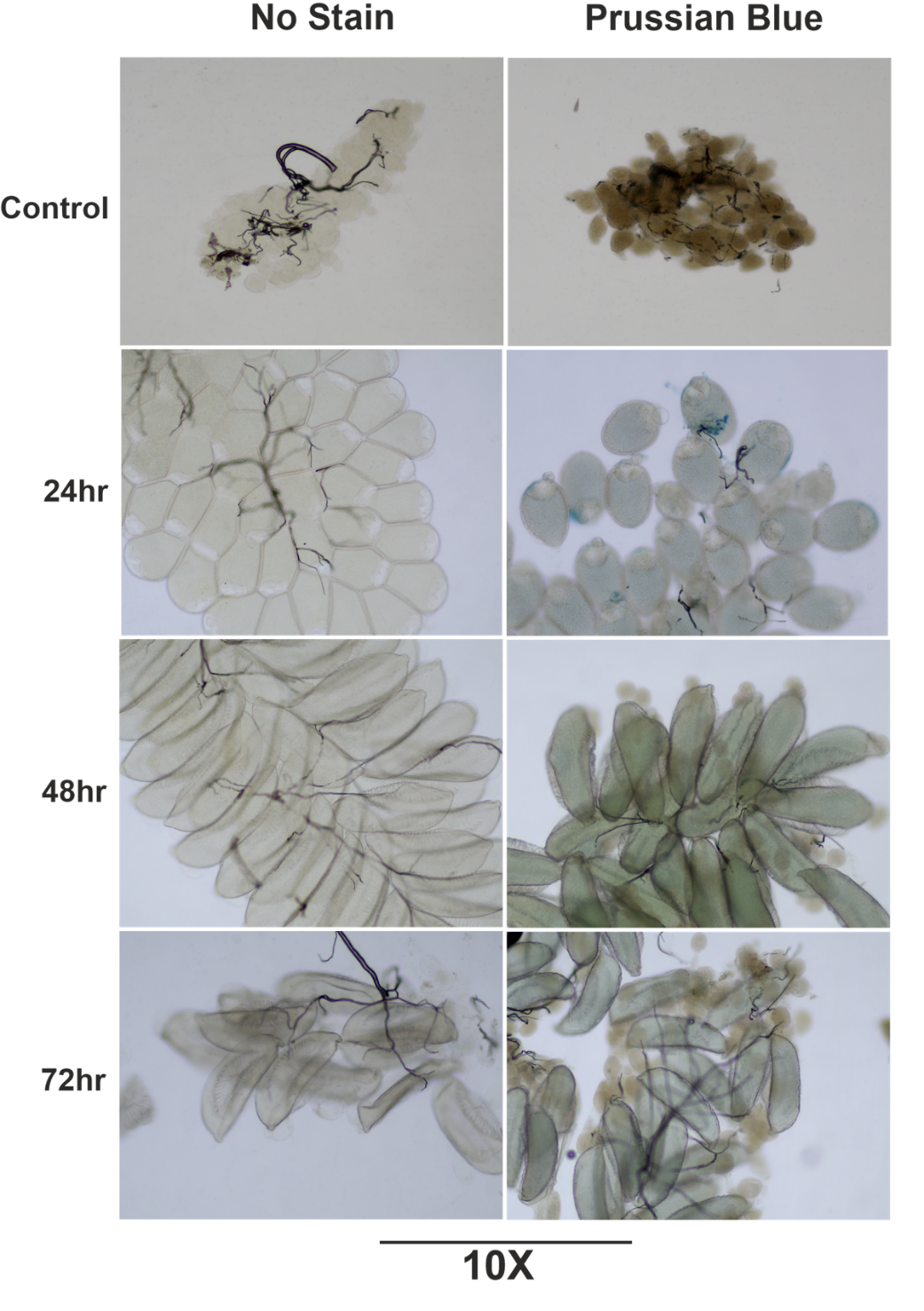
**Figure S6.** **Prussian blue staining reveals temporal accumulation of iron in *Anopheles stephensi* ovaries following a blood meal.** Ovaries were dissected from control (unfed) females and from females at 24-, 48-, and 72-hours post-blood meal (PBM), and imaged either without stain (left column) or after Prussian blue staining (right column) to visualize ferric iron deposits. Iron accumulation becomes increasingly prominent in ovarian tissues at 48, and 72 hours PBM, as indicated by the progressive intensification of blue coloration. Images were captured and visualised at 10× magnification under Olympus BX63 Light Microscope.

**C**

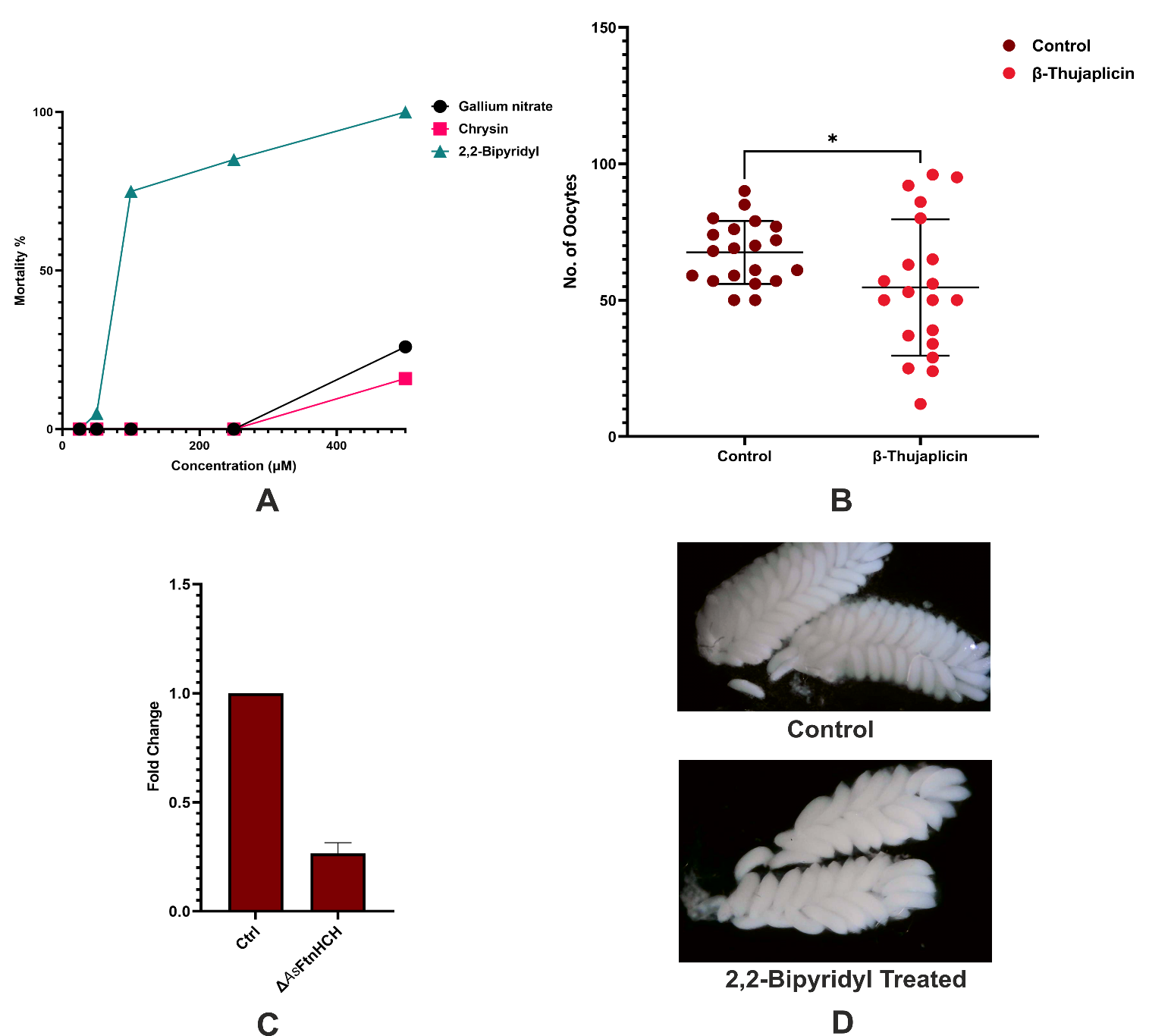

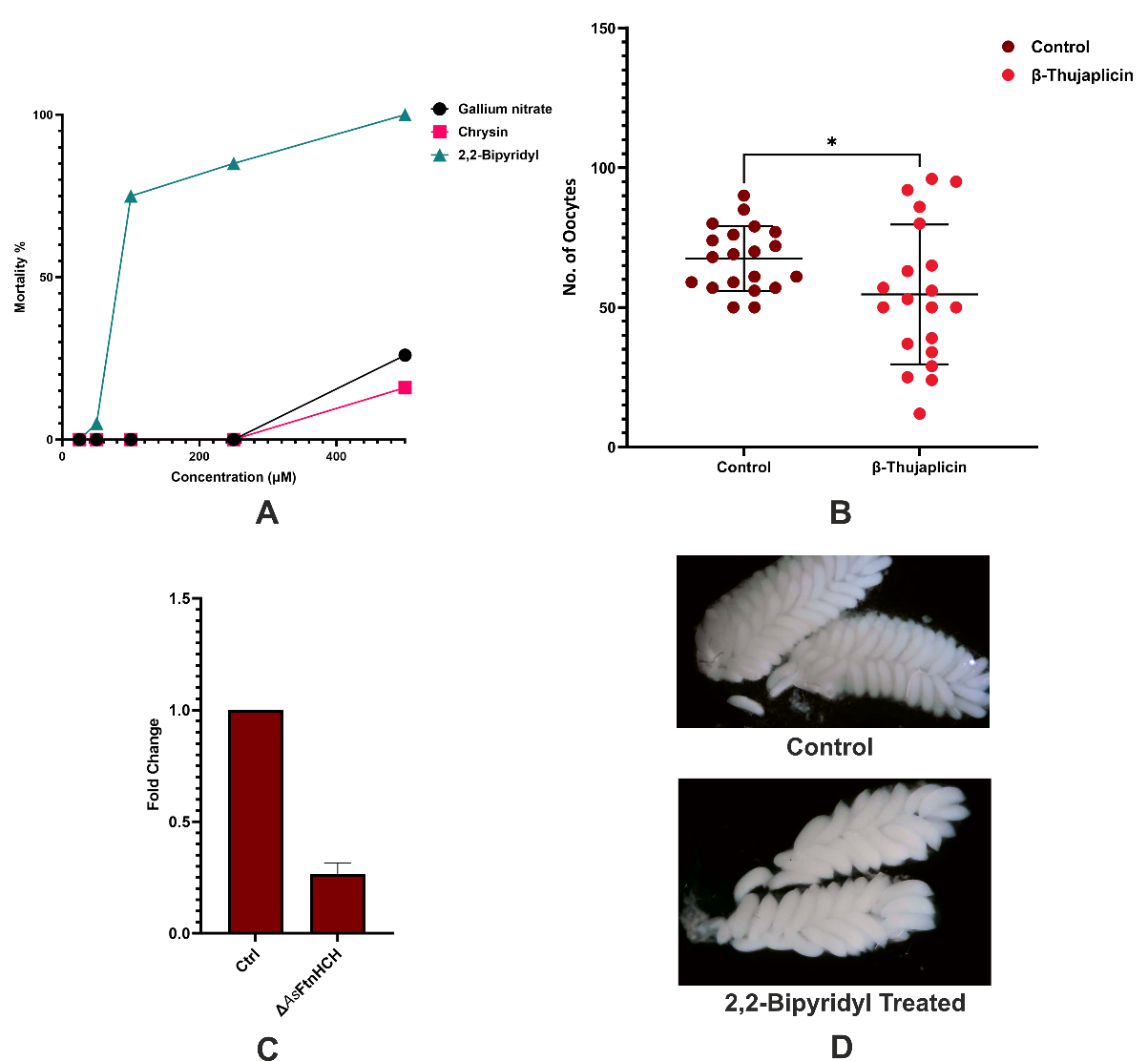

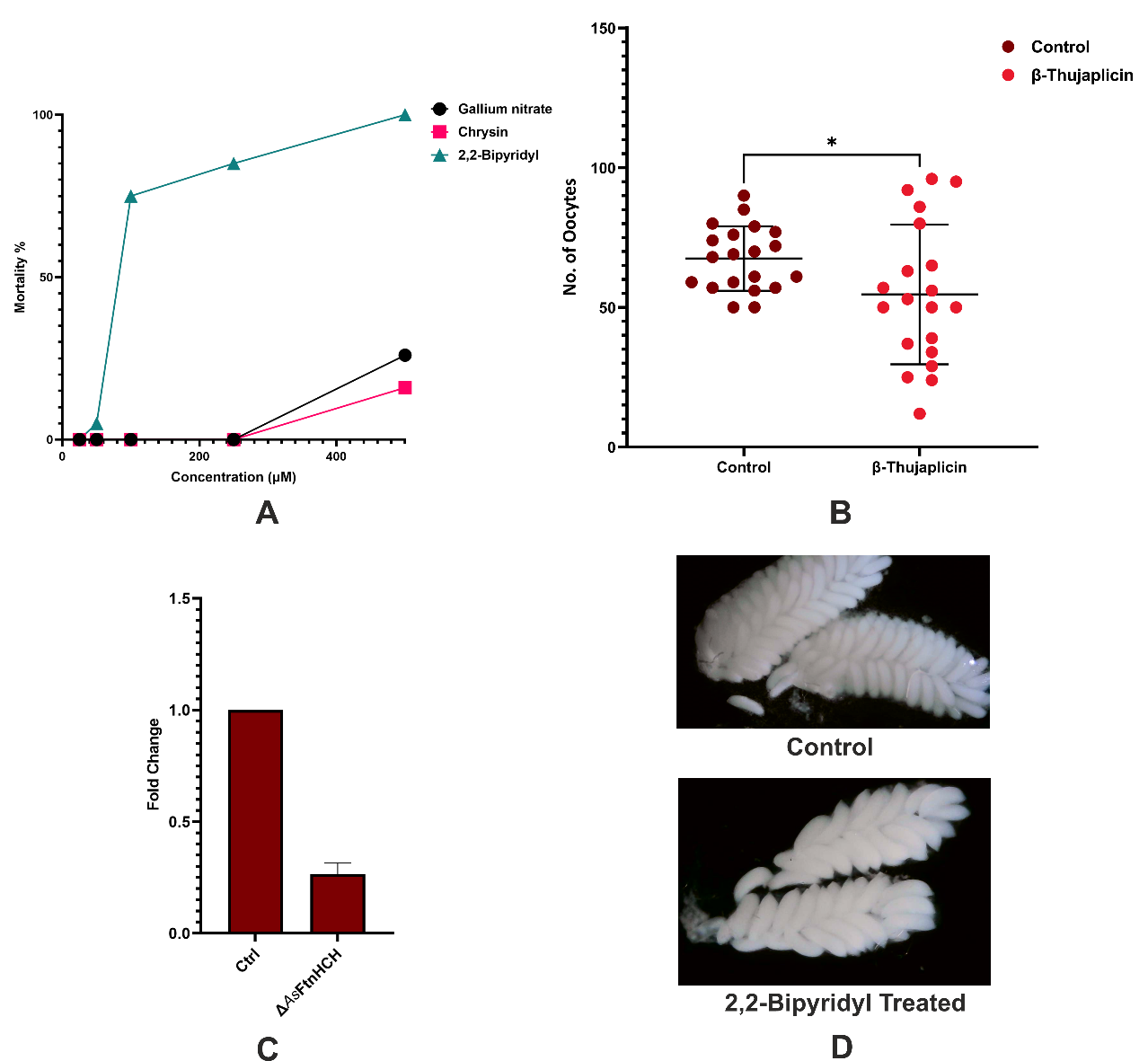

**A**

**B**

**Figure S7.** **Effect of iron chelation on the development of *Anopheles stephensi*.** (A) Dose-dependent larval mortality (%) in *Anopheles stephensi* upon iron chelator treatment (n=30); (B) Microscopic images of dissected ovaries from control and 2,2′-bipyridyl-treated *Anopheles stephensi* females, illustrating the marked reduction in oocyte development upon chelator treatment.; (C) Quantification of oocyte numbers per female in control and β-Thujaplicin-treated groups, showing reduction in oocyte numbers following iron chelation (*p < 0.05); each dot represents an individual mosquito, with bars indicating mean ± SD (n = 20).;

**Table S1.** List of conserved iron-binding residues in Human heavy chain ferritin, *Anopheles culicifacies*’s ferritin

heavy chain homolog and bacterioferritin from *Desulfovibrio desulfuricans*.

| Human ferritin | Mosquito ferritin  (*An.culicifacies*) | Bacterioferritin  (*Desulfovibrio desulfuricans*) |
| --- | --- | --- |
| Glu28 | Glu58 | Glu23 |
| Glu63 | Glu93 | Glu56 |
| His66 | His96 | His59 |
| Glu108 | Glu147 | Glu98 |
| Gln148 | Gln182 | Glu132 |
| - | - | His135 |

**Table S2.** Chemical and structural profiles of iron chelators used in the study.

| Name | Mol. weight | logP Value* | Chemical Structure |
| --- | --- | --- | --- |
| 2,2-Bipyridyl | 156.18 g/mol | 2.94 | 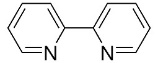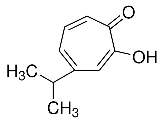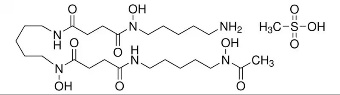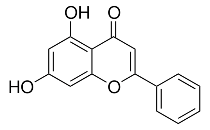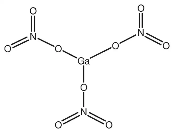 |
| β-Thujaplicin | 164.20 g/mol | 1.68 |  |
| Deferoxamine Mesylate | 656.79 g/mol | 0.614 |  |
| Chrysin | 254.24 g/mol | 2.68 |  |
| Gallium Nitrate | 255.74 g/mol | -3.05 |  |

*Negative values indicate hydrophilic property

**Table S3**. List of all the primers used in the study.

| Name | Forward primer (5'-3') | Reverse Primer (5'-3') |
| --- | --- | --- |
| *As*FtnHCH | ATTGACCGCAACACCTTCTC | CTTAATTAGCTCGCGGATGC |
| S7 | AGCTGGAGCGGTACCATTTG | CCACGCTTGATCGGTGTC |

**Table S4.** Amino acid sequence of ferritin’s synthetic construct used in the study.

| **Species name** | **Amino acid sequence** |
| --- | --- |
| *Homo sapiens* (Hftn) | MTTASTSQVRQNYHQDSEAAINRQINLELYASYVYLSM  SYYFDRDDVALKNFAKYFLHQSHEEREHAEKLMKLQN  QRGGRIFLQDIKKPDCDDWESGLNAMECALHLEKNVN  QSLLELHKLATDKNDPHLCDFIETHYLNEQVKAIKELG  DHVTNLRKMGAPESGLAEYLFDKHTLGDSDNES |
| *Anopheles culicifacies* (*Ac*FtnHCH) | MQVTDTDSPSGVDEWNYMNRSCSAKLQDQINKEFDA  AIFYMQYAAYFAQYKVNLPGFEKFFFEAASEEREHGM  KLIEYALMRGQKPIDRNSFSLHFANQVQQPEPEQGSVA  LTALRAALAKEQEVTKSIRELIKICEEDHNDYHLVDYLT  GEFLEEQHQGQRDLAGKITMLAKLLRTNPKLGEFMFDK  QNM |
| *Acinetobacter baumannii* (Bftn) | MKGNRDVINQLNQVLYHHLTAINQYFLHSRMFNDWGI  EQLGSAEYKESIRQMKHADKIIERILFLEGLPNLQHLGK  LYIGQHTEEVLQCDIRKVKENIEAIQKAVALAETEQDYV  TRDLVQEILEKEEEYWDWLDTQIDLIGSVGIENYIQSQM |

**Table S5**. List of all the species and their UniProt ID used to generate the phylogenetic tree.

| **Figure S1B***  **Species name**  **(UniProtID)** | | **Figure S1A***  **Species name**  **(UniProtID)** | | **Figure 1B***  **Species name (UniProtID)** |
| --- | --- | --- | --- | --- |
| *Anopheles culicifacies*  (A0A182M0V3) | *Aedes aegypti*  (P41822) | *Anopheles gambiae*  (Q7QC60) | *Hyalomma anatolicum*  (A0A0D5W3A6) | *Anopheles culicifacies*  (A0A182M0V3) |
| *Anopheles minimus*  (A0A182VXD7) | *Anopheles sinensis*  (A0A084WRD4) | *Aedes aegypti*  (P41822) | *Haemaphysalis longicornis*  (Q6WNX1) | *Aedes aegypti*  (P41822) |
| *Anopheles funestus*  (A0A182RA08) | *Anopheles melas*  (A0A182U8R7) | *Apis mellifera*  (A0A7M7MU23) | *Haemaphysalis flava*  (A0A8E4NKH1) | *Culex quinquefasciatus*  (B0X505) |
| *Anopheles stephensi*  (A0A182XWX0) | *Culex quinquefasciatus*  (B0X505) | *Bemisia tabaci*  (A0A2U9QWK6) | *Hyalomma rufipes*  (UPI0021FF7335) | *Gallus gallus*  (P08267) |
| *Anopheles christyi*  (A0A182JRI0) |  | *Asobara tabida*  (D1GLP2) | *Manduca sexta*  (Q8WQX3) | *Homo sapiens*  (P02794) |
| *Anopheles quadriannulatus*  (A0A182XI13) |  | *Apriona germarii*  (Q8MUX0) | *Leptinotarsa decemlineata*  (Q7Z161) | *Macaca mulatta*  (F7F2L7) |
| Anopheles arabiensis  (A0A182HSZ6) |  | *Bactrocera dorsalis*  (A0A059XRL4) | *Nilaparvata lugens*  (Q9U0S2) | *Mus musculus*  (P09528) |
| *Anopheles gambiae*  (Q7QC60) |  | *Bombus ignites*  (A8D916) | *Musca domestica*  (T1PLJ3) | *Bos taurus*  (O46414) |
| *Anopheles merus*  (A0A182V5L6) |  | *Bombyx mori*  (Q1HQ56) | *Ornithodoros moubata*  (O61916) | *Xenopus laevis* (P49948) |
| *Anopheles epiroticus*  (A0A182PAE6) |  | *Dermacentor variabilis*  (Q8T9S6) | *Ixodes Ricinus*  (A0A0K8RA47) | *Danio rerio*  *(*Q9DDT0) |
| *Anopheles atroparvus*  (A0A182JMC7) |  | *Chilo suppressalis*  (A0ABN8B5B7) | *Papilio Xuthus*  (I4DJ24) | *Caenorhabditis elegans*  (Q9TYS3) |
| *Anopheles farauti*  (A0A182Q958) |  | *Camponotus floridanus*  (E2AZU0) | *Trichoplusia ni*  (A0A7E5WTY7) | *Arabidopsis thaliana*  (Q9S756 |
| *Anopheles maculatus*  (A0A182SSI9) |  | *Galleria mellonella*  (Q9GQW5) | *Samia ricini*  (A0A2D1AEE0) | *Escherichia coli*  (P0ABD3) |
| *Anopheles albimanus*  (A0A182FT42) |  | *Glossina morsitans* (Q2PYZ6) | *Rhipicephalus sanguineus*  (Q6WNW9) | *Cyanobacterium aponinum*  (A0AAF0ZA07) |
| *Anopheles darlingi*  (W5JML5) |  | *Drosophila melanogaste*r  (Q7KRU8) | *Rhodnius prolixus*  (R4G2V0) | *Thermotoga maritima*  (Q9WZ30) |

*Respective phylogentic tree
